# Deletion of calcineurin from astrocytes reproduces proteome signature of Alzheimer’s disease and epilepsy and predisposes to seizures

**DOI:** 10.1101/2020.03.21.001321

**Authors:** Laura Tapella, Giulia Dematteis, Federico Alessandro Ruffinatti, Luisa Ponzoni, Fabio Fiordaliso, Alessandro Corbelli, Enrico Albanese, Beatrice Pistolato, Jessica Pagano, Elettra Barberis, Emilio Marengo, Claudia Balducci, Gianluigi Forloni, Chiara Verpelli, Carlo Sala, Carla Distasi, Mariaelvina Sala, Armando A. Genazzani, Marcello Manfredi, Dmitry Lim

**Author notes:** LT and GD contributed equally to this work. These Authors share seniorship (Marcello Manfredi, Dmitry Lim). Correspondence should be send to Dmitry Lim.

## Abstract

In astrocytes, calcineurin (CaN) is involved in neuroinflammation and gliosis, while its role in healthy CNS or in early neuro-pathogenesis is poorly understood. Here we report that in astroglial CaN KO (ACN-KO) mice, at one month of age, proteome is deranged in hippocampus and cerebellum. Bioinformatic analysis reveals association with Alzheimer’s disease (AD) and epilepsy. We found significant overlap with the proteome of an AD mouse model and of human subjects with drug-resistant epilepsy. In Barnes maze ACN-KO mice learned the task but adopted serial search strategy. Strikingly, from five months of age ACN-KO mice develop spontaneous seizures with an inflammatory signature of epileptic brains. These results suggest that astroglial CaN KO impairs hippocampal connectivity, produces proteome features of neurological disorders and predisposes mice to seizures. We suggest that astroglial CaN may serve as a novel Ca2+-sensitive switch which regulates protein expression and homeostasis in the CNS.

## INTRODUCTION

Homeostatic support, provided by astrocytes, is of paramount importance for proper brain functioning (Verkhratsky & Nedergaard, 2018). During continuous interaction with other cells in the central nervous system (CNS), astrocytes generate intracellular calcium (Ca^2+^) signals which are thought to mediate their homeostatic activities, including release of signalling molecules, control of cerebral blood flow, regulation of K^+^ and neurotransmitter uptake and control of neuronal activity (Khakh & Sofroniew, 2015). Moreover, deregulation of astrocytic Ca^2+^ signals and downstream signalling processes are strongly associated with neuropathologies including Alzheimer’s disease and epilepsy (Lim, Rodríguez-Arellano, et al., 2016; Nikolic et al., 2019). Much effort has been devoted to detection and characterization of astroglial Ca^2+^ signals themselves (Bindocci et al., 2017; Fiacco et al., 2009), while how the Ca^2+^ signals in astrocytes are decoded, processed and translated to arrange a multitude of astroglial homeostatic activities is poorly understood.

Recently, we have generated a mouse with conditional KO of calcineurin (CaN), a Ca^2+^/calmodulin-activated serine-threonine phosphatase (<u>a</u>stroglial <u>c</u>alci<u>n</u>eurin <u>KO</u>, or ACN-KO) and showed that the deletion of astrocytic CaN (Astro-CaN) severely impairs intrinsic neuronal excitability through functional inactivation of astroglial Na^+^/K^+^ ATPase and deregulation of Na^+^ and K^+^ handling (Tapella et al., 2020). CaN drives Ca^2+^-dependent cellular transcriptional and functional remodelling in different cellular types including lymphocytes (Oh-hora & Rao, 2008), cardiomyocytes (Parra & Rothermel, 2017), skeletal muscle (Tu et al., 2016), bone cells (Sitara & Aliprantis, 2010) and neurons (Baumgärtel & Mansuy, 2012). However, it is not known whether in healthy astrocytes CaN exerts similar activity.

In the present report we used a multiomics approach to comprehensively investigate alterations produced by the deletion of Astro-CaN at the transcriptional and proteome level. We report that the deletion of CaN from astrocytes, at one month of age, results in derangement of protein expression at the posttranscriptional level re-creating a similar protein profile to that associated with neurological and neurodegenerative diseases, in particular with AD and seizures. Later in life, beginning from about 5 months of age, ACN-KO mice display an increased risk to develop spontaneous tonic-clonic motor seizures. Collectively, our results suggest that Astro-CaN may serve as a Ca^2+^-sensitive molecular hub which regulates protein expression and homeostasis in the CNS.

## RESULTS

### Deletion of Astro-CaN deranges protein expression at posttranscriptional level

To investigate the effect of the deletion of CaN from astrocytes on gene and protein expression in the CNS, we performed Next Generation RNA sequencing (RNA-Seq) and shotgun mass spectrometry proteomics (SG-MS) analyses of the hippocampus and cerebellum of one month-old ACN-KO and ACN-Ctr mice. RNA-Seq releaved only two genes that showed a fold change over 2 in whole tissue hippocampal and cerebellar preparations from ACN-KO compared with ACN-Ctr mice (Supplementary Figure 1). This were Rab31 and Vapa in the hippocampus (−2.11 and −2.04 fold change, respectively) and Dsp and Rab31 in cerebellum (−2.62 and −2.14 fold change, respectively). In brief, therefore, the transcriptional profile was largely unchanged. SG-MS proteomics (Fig. 1A) identified 1661 and 1456 proteins in whole-tissue hippocampal (Wht-Hip) and cerebellar (Wht-Cb) preparations, respectively. In hippocampal (Syn-Hip) and cerebellar (Syn-Cb) synaptosomal fractions 525 and 471 proteins, respectively, were identified (Supplementary Table 1). Quantification and identification of differentially expressed proteins (DEPs) (FC > 50%, p-value < 0.05) yielded 21, 17, 134 and 40 DEPs in Wht-Hip, Wht-Cb, Syn-Hip and Syn-Cb samples, respectively (Fig. 1B and Supplementary Table 2). Surprisingly, there was very little, if any, overlap of DEPs between samples suggesting that the changes in protein expression produced by the deletion of Astro-CaN are brain region-specific. Validation of SG-MS results by WB (Fig. 1C) showed significant correlation (R^2^ = 0.612, p = 0.0044) between SG-MS and WB (Fig. 1D).

**Figure 1.**
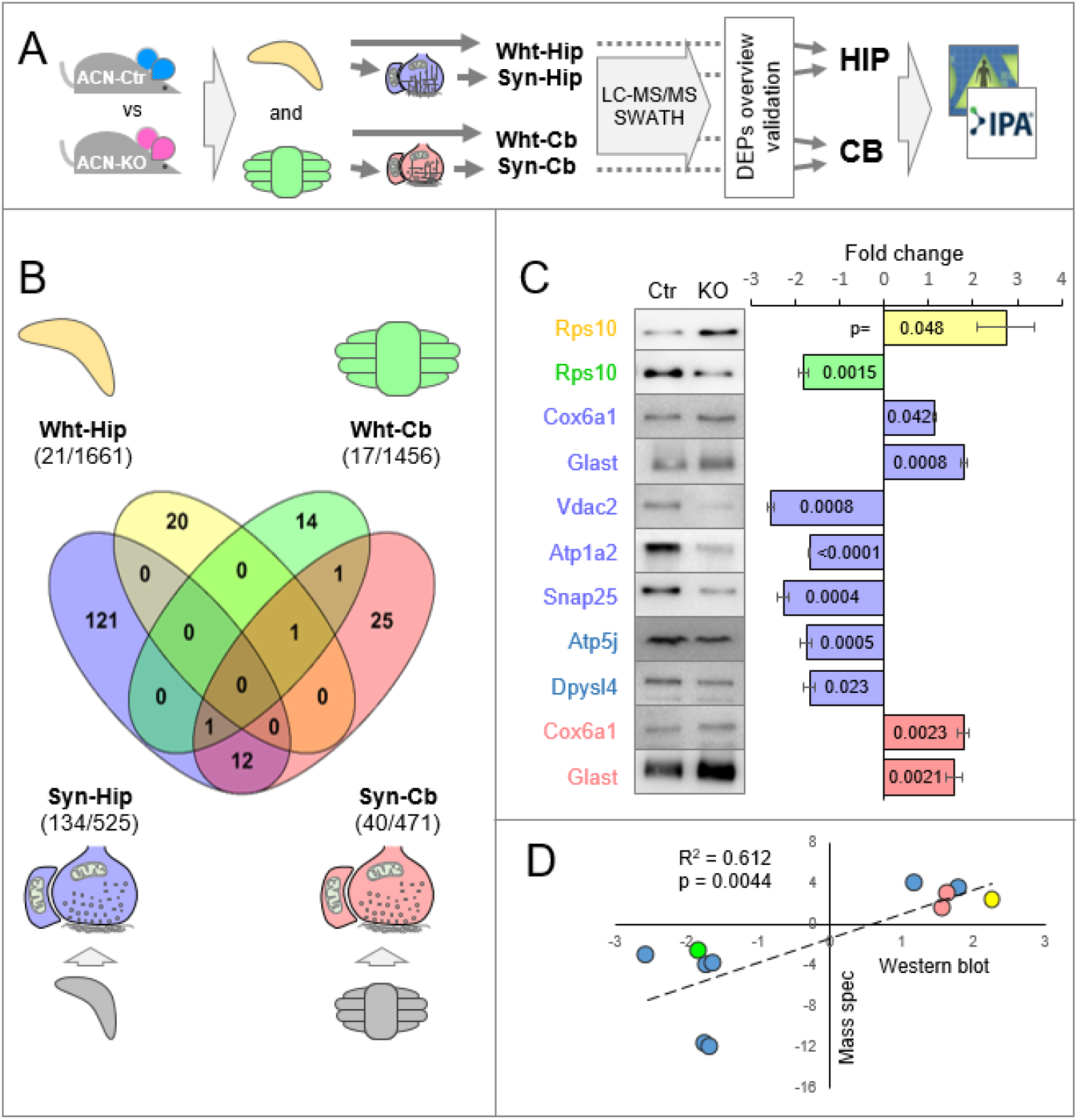
Astrocyte-specific CaN knockout at 1 mo of age deranges protein expression differentially in the hippocampus and the cerebellum. (A) A scheme of the workflow: whole-tissue hippocampus (Wht-Hip) and cerebellum (Wht-Cb) and their respective synaptosomal fractions (Syn-Hip and Syn-Cb), prepared from ACN-Ctr and ACN-KO mice, were subjected either to NGS RNA sequencing (RNA-Seq, n = 6 mice per condition) or to shotgun mass spectrometry (SG-MS) proteomics (LC-MS/MS – SWATH, 4-6 mice per condition). RNA-Seq did not result in a significant transcriptional activity. SG-MS resulted in 21, 17, 134 and 40 differentially expressed proteins (DEPs) in Wht-Hip, Wht-Cb, Syn-Hip and Syn-Cb, respectively. For bioinformatic analysis, hippocampal and cerebellar preparations were pooled resulting in HIP (155 DEPs) and CB (54 DEPs). (B) Intersection of DEPs lists show very limited overlap between samples indicating on brain region-specific alterations produced by Astro-CaN-KO. (C) Validation of DEPs by Western blot; and (D) correlation of Western blot results with SG-MS (R^2^ = 0.612, p = 0.0044). Colour codes: cyan, ACN-Ctr mice; magenta, ACN-KO mice; light-yellow, Wht-Hip, light-green, Wht-Cb; lightblue, Syn-Hip; light-red, Syn-Cb.

qPCR validation showed no alterations in mRNA levels except Vapa and Rab31, found to be downregulated in hippocampus confirming, thus, RNA-Seq results (Supplementary Figure 2). Classification of DEPs for Cell component and Function categories revealed that in Wht-Hip and Wht-Cb Cell components of *cytosol* were the most abundant followed by *mitochondria, ribosome* and *vesicles,* while among Functions there were *metabolism, translation, OXPHOS* followed by *vesicle trafficking* and *transport across membrane.* In synaptosomal preparations Cell components were categorized as *cytosol, cytoskeleton, mitochondria, plasma membrane, vesicles* and *ribosome;* while among Functions were *signalling, metabolism, transport across membrane, structural proteins, OXPHOS, translation* and *vesicle trafficking* (Supplementary Figure 3).

### Deletion of Astro-CaN deranges protein expression in major brain cell types

The analysis of DEPs showed the presence of many neuronal proteins, e.g., Thy-1, Snap25, Synaptophysin and Tau suggesting that the deletion of CaN from astrocytes affects neuronal protein expression. Therefore, we analysed the cell-specificity of DEPs using a merged lists of Wht-Hip and Syn-Hip DEPs (Supplementary Figure 2E). First, we primed the Human Protein Atlas (HPA) database (https://www.proteinatlas.org/) in which protein expression is comprehensively examined using ATLAS antibodies (Sigma) and the cell-specificity of expression in the hippocampal formation is scored in glial and neuronal cells. This analysis, revealed that, in the hippocampus, the majority of DEPs were neuron-specific (32.9%) or enriched in neurons (24.5%), while glia-specific proteins (6.5%) or proteins enriched in glia (2.6%) were less abundant (Fig. 2A). Next, we primed a brain cell-specific transcriptome database published by Ben Barres and colleagues (Zhang et al., 2014). The analysis revealed that DEPs specific for all principal brain cell types, including astrocytes (8 hits), neurons (19 hits), oligodendrocytes (7 hits), microglial cells (3 hits) and endothelial cells (2 hits) were differentially expressed in ACN-KO brain (Fig. 2B), corroborating and extending the results obtained using HPA.

**Figure 2.**
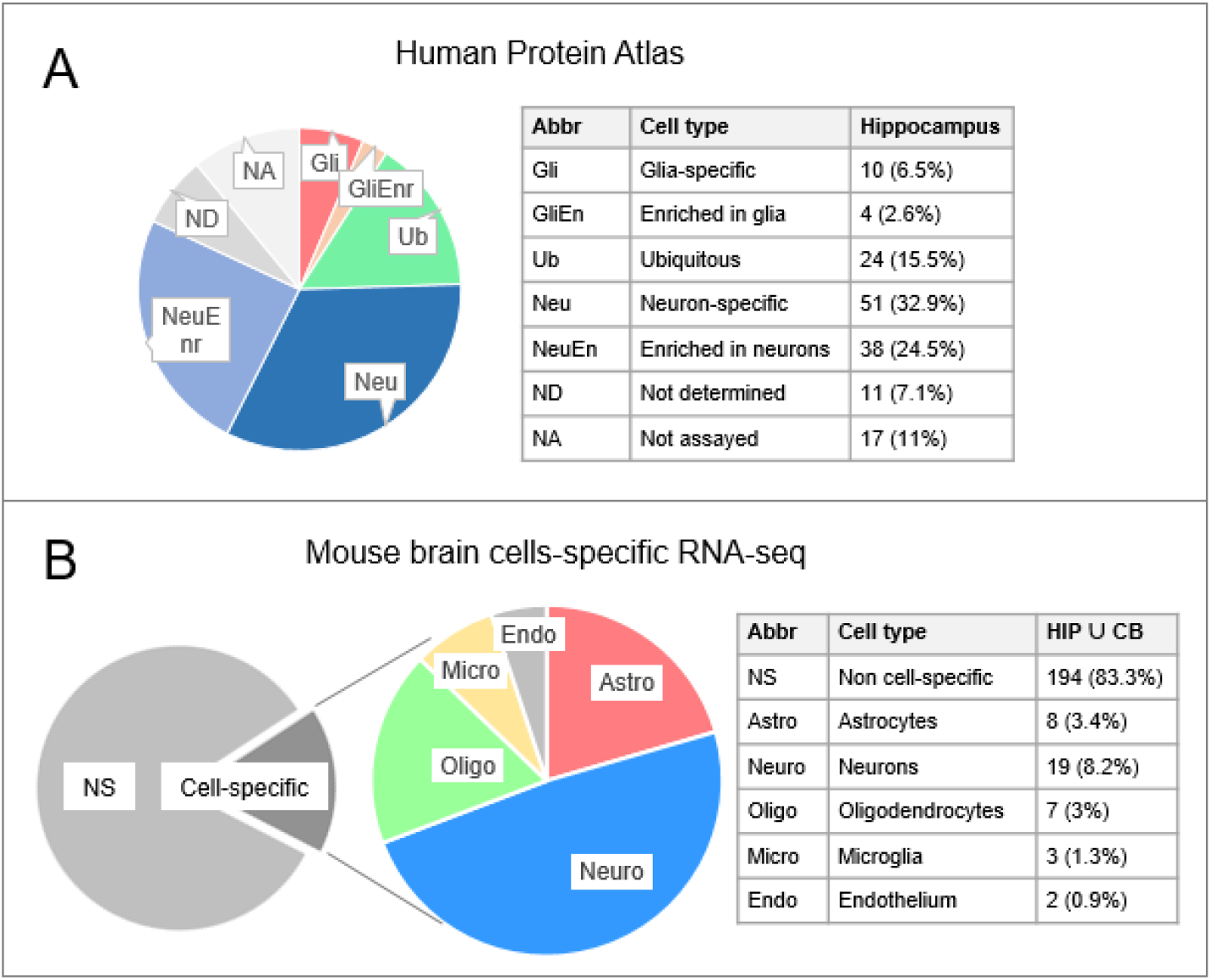
Cell-specificity analysis of differentially expressed proteins. (A) Priming of Human Protein Atlas database (www.proteinatlas.org) of Hippocampal differentially expressed proteins (DEPs) revealed prevalence of neuronal DEPs. (B) Priming of mouse brain cell-specific RNA-Seq transcriptome revealed alterations in expression of proteins from all major CNS cell types, with prevalence of neuronal proteins.

Taken together, RNA-Seq and SG-MS analyses showed that at one month of age the deletion of Astro-CaN does not change transcription neither in the hippocampus nor in the cerebellum, while protein expression was altered differentially in these two brain regions, in all principal brain cellular types.

### Deletion of Astro-CaN produces proteome signature of neuropathology and seizures

Next, we investigated the functional significance of protein expression alterations in ACN-KO mice. For this analysis, whole-tissue DEPs were pooled with synaptosomal DEPs resulting in two lists, related to the hippocampus (HIP) and the cerebellum (CB). Gene ontology analysis of HIP DEPs revealed overrepresentation of 175 GO terms (Supplementary Table 3A). The most significantly overrepresented hits from GO category Cellular Component (CC) were *myelin sheath* (44 DEPs), *extracellular exosome* (90 DEPs) and *mitochondrion* (46 DEPs), while KEGG pathways were *OXPHOS* (17 DEPs), *Alzheimer’s* (AD, 13 DEPs), *Parkinson’s* (PD, 15 DEPs) and *Hunlinglon’s disease* (HD, 15 DEPs). Interestingly, intersection of *AD, PD, HD* and *OXPHOS* lists resulted in 12 common DEPs, all of which were components of the oxidative phosphorylation (OXPHOS) mitochondrial pathway (Supplementary Figure 4). DAVID analysis of CB list returned 43 significantly overrepresented GO terms among which were KEGG pathway *ribosome* (7 DEPs) and CC *extracellular exosome* (33 DEPs), *intermediate filament* (7), *intracellular ribonucleoprotein complex* (9), *ribosome* (7) and *mitochondrion* (15) (Supplementary Table 3B). Presence of intermediate filaments of the keratin family was somewhat surprising, however priming databases Genecards (https://www.genecards.org/) and PaxDB (https://pax-db.org/) revealed presence of keratins in different brain proteome datasets (Supplementary Table 4).

Ingenuity pathway analysis (IPA) of HIP dataset resulted in a list of at least 500 significantly overrepresented annotations. Top Canonical Pathways were *Synaptogenesis Signaling Pathway, Phagosome Maturation, Sirtuin Signaling Pathway,* and *Mitochondrial Dysfunction* (Table 1). Overrepresented Disease And Functions annotations contained developmental and hereditary neurological disorders as well as psychiatric and neurodegenerative disorders (full list of annotations provided in Supplementary Table 5). Note the presence of annotations related to seizures and epilepsy (Table 1) which contained 29 proteins (listed in Supplementary Table 6). Top Upstream Regulators in HIP were proteins *MAPT, PSEN1, APP* and *HTT* (Table 1 and Supplementary Table 7A). Mutations of MAPT (tau), PSEN1 (presenilin 1) and APP (amyloid precursor protein) are the major causes of familial AD (FAD) (Loy et al., 2014), while mutation of HTT is a cause of Huntington’s disease (Nance, 2017). The relationships of MAPT, PSEN1 and APP with target proteins, calculated by IPA, are shown, respectively, in Supplementary Figures 5, 6 and 7. Of 69 unique proteins in *MAPT, PS1, APP* lists, 47 were common (Supplementary Table 7B), falling in three clusters, generated using STRING unsupervised k-means clustering, (i) neuronal and synaptic, (ii) metabolic pathways, mitochondrial and OXPHOS, and (iii) cytoskeletal and adhesion proteins (Fig. 3, and Supplementary Table 7C). Top Canonical Pathways in CB list were *EIF2 Signaling, Glucocorticoid Receptor Signaling, Regulation of eIF4 and p70S6K Signaling, Sirtuin Signaling Pathway* and *Mitochondrial Dysfunction.* Top Upstream Regulators in CB were *MAPT, MYC* and *RICTOR,* while Disease And Functions annotations included *Decay of mRNA, Synthesis of protein, Parkinson’s disease* and *Progressive encephalopathy* (Table 1).

**Figure 3.**
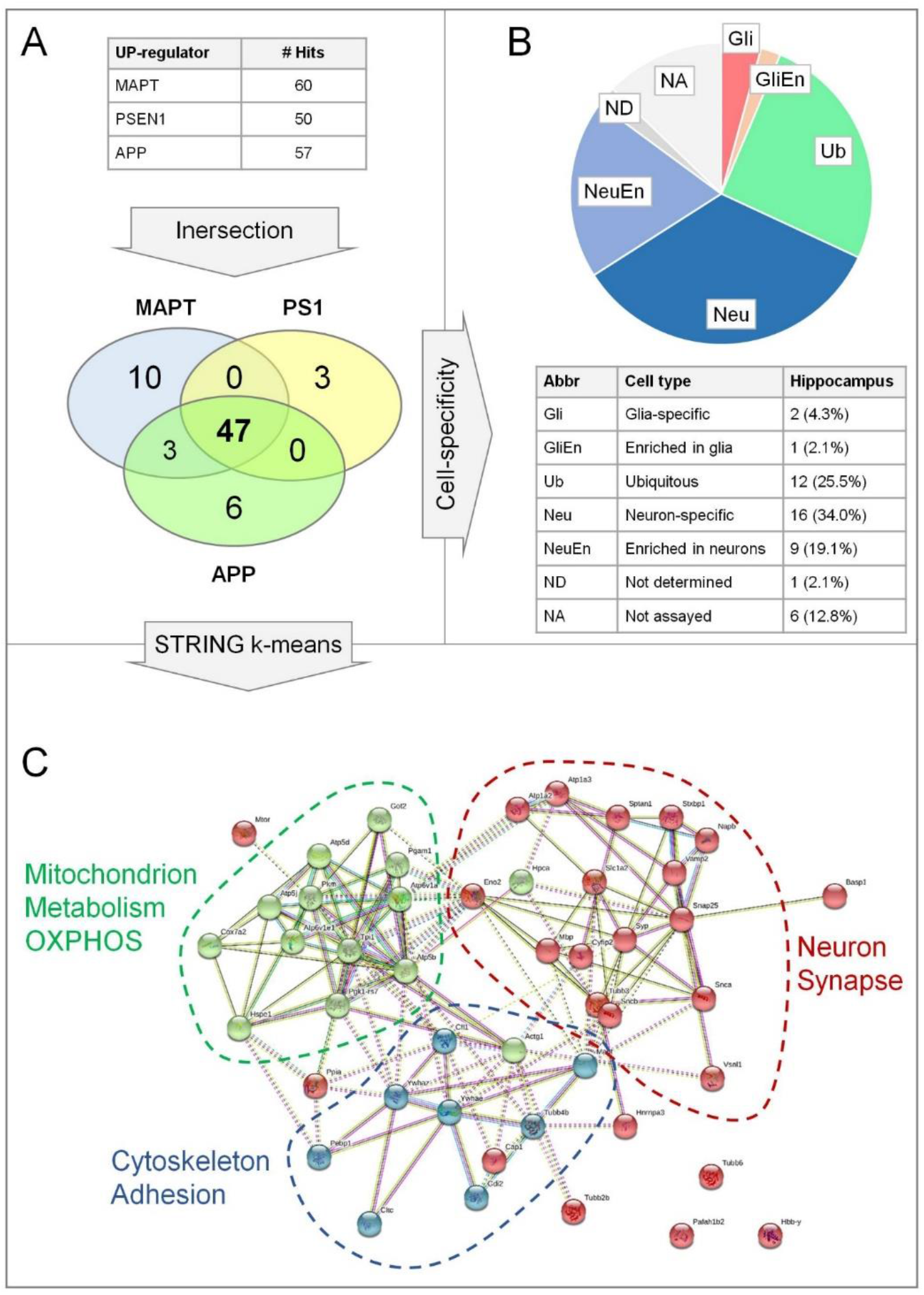
Top 3 Cell-specificity and STRING k-means clustering of common targets of IPA pathway Upstream Regulators of Hippocampal DEPs MAPT, PS1 and APP. (A) Intersection of 60 MAPT, 50 PS1 and 57 APP putative regulated DEPs resulted in 47 common DEPs. (B) Cell-specificity analysis Huma Protein Atlas (www.proteinatlas.org) database shows presence of mostly neuronal and neuron-enriched DEPs. (C) 47 common DEPs regulated by MAPT, PS1 and APP, putative upstream regulators found by IPA pathway analysis, were subjected to STRING proteinprotein interaction analysis. STRING k-means unsupervised clustering revealed 3 clusters grouped in proteins related to: neuron and synapse (Cluster 1, red), mitochondrion, metabolism and OXPHOS (Cluster 2, green); and cytoskeleton and adhesion (Cluster 3, light-blue).

**Table 1.**
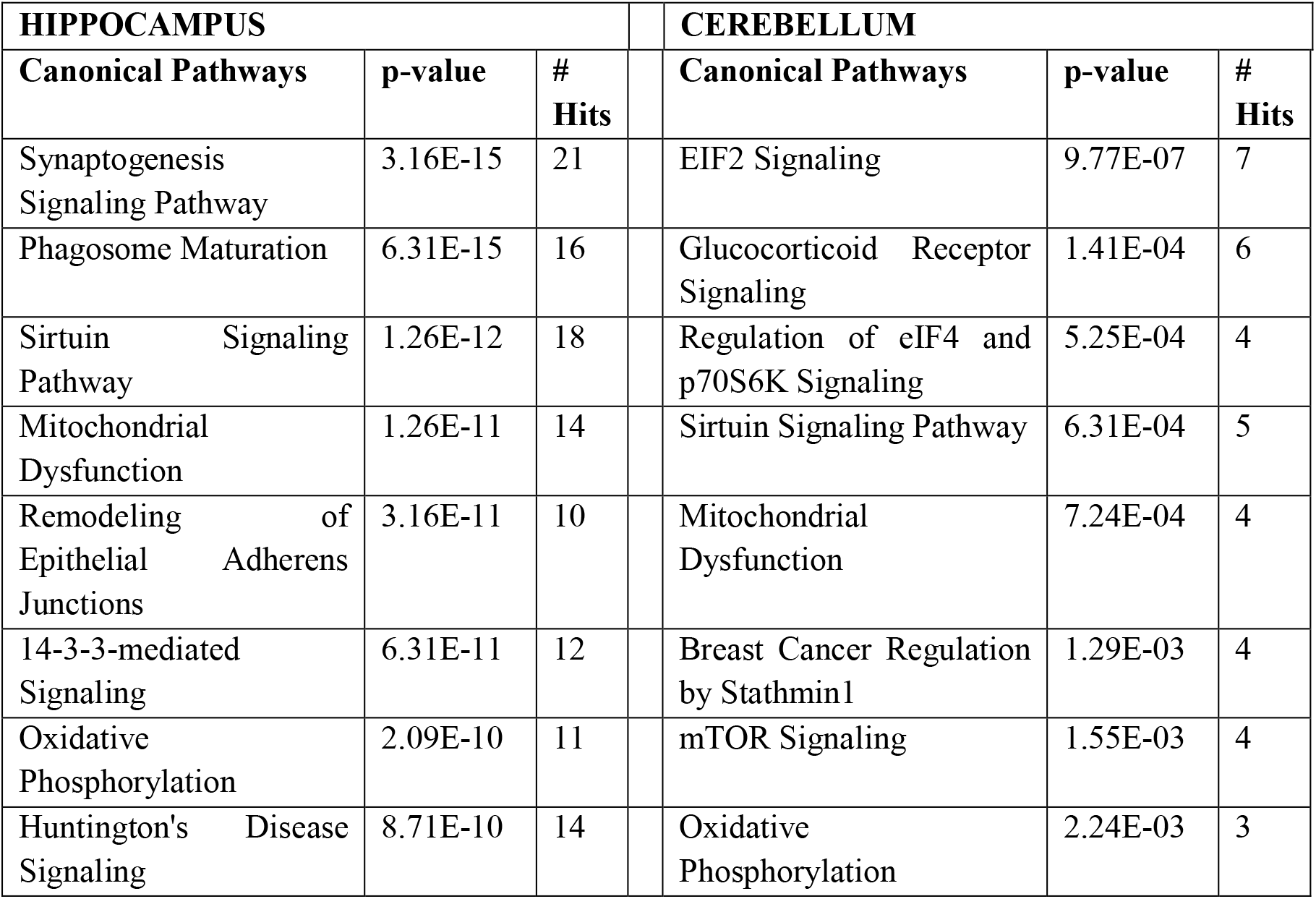
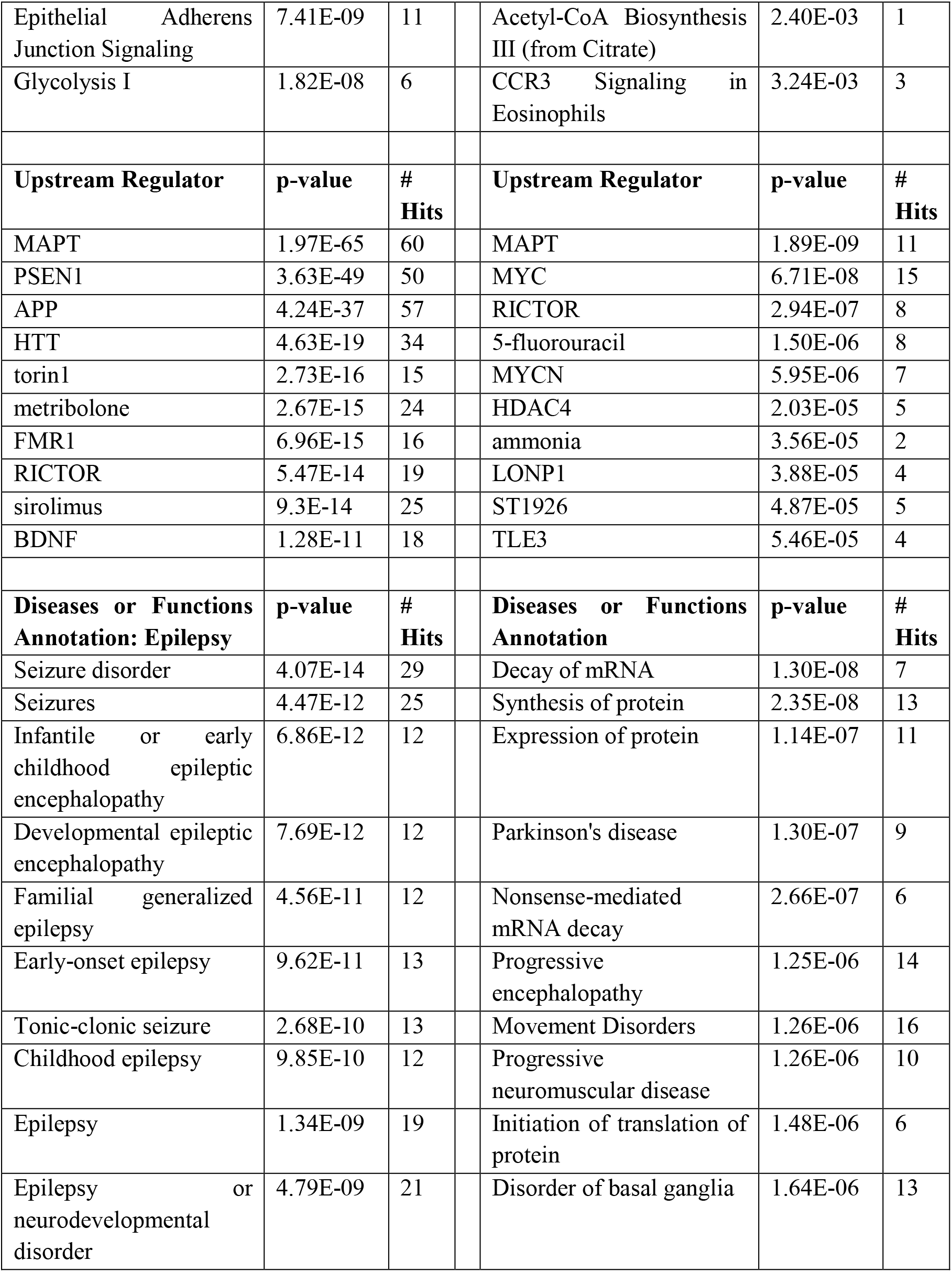
Top 10 overrepresented IPA annotations.

| HIPPOCAMPUS |  |  | CEREBELLUM |  |  |
| --- | --- | --- | --- | --- | --- |
| Canonical Pathways | p-value | # Hits | Canonical Pathways | p-value | # Hits |
| Synaptogenesis Signaling Pathway | 3.16E-15 | 21 | EIF2 Signaling | 9.77E-07 | 7 |
| Phagosome Maturation | 6.31E-15 | 16 | Glucocorticoid Receptor Signaling | 1.41E-04 | 6 |
| Sirtuin Signaling Pathway | 1.26E-12 | 18 | Regulation of eIF4 and p70S6K Signaling | 5.25E-04 | 4 |
| Mitochondrial Dysfunction | 1.26E-11 | 14 | Sirtuin Signaling Pathway | 6.31E-04 | 5 |
| Remodeling of Epithelial Adherens Junctions | 3.16E-11 | 10 | Mitochondrial Dysfunction | 7.24E-04 | 4 |
| 14-3-3-mediated Signaling | 6.31E-11 | 12 | Breast Cancer Regulation by Stathmin1 | 1.29E-03 | 4 |
| Oxidative Phosphorylation | 2.09E-10 | 11 | mTOR Signaling | 1.55E-03 | 4 |
| Huntington's Disease Signaling | 8.71E-10 | 14 | Oxidative Phosphorylation | 2.24E-03 | 3 |

|  |  |  |  |  |  |  |
| --- | --- | --- | --- | --- | --- | --- |
| Epithelial Adherens Junction Signaling | 7.41E-09 | 11 |  | Acetyl-CoA Biosynthesis III (from Citrate) | 2.40E-03 | 1 |
| Glycolysis I | 1.82E-08 | 6 |  | CCR3 Signaling in Eosinophils | 3.24E-03 | 3 |
| **Upstream Regulator** | **p-value** | **# Hits** |  | **Upstream Regulator** | **p-value** | **# Hits** |
| MAPT | 1.97E-65 | 60 |  | MAPT | 1.89E-09 | 11 |
| PSEN1 | 3.63E-49 | 50 |  | MYC | 6.71E-08 | 15 |
| APP | 4.24E-37 | 57 |  | RICTOR | 2.94E-07 | 8 |
| HTT | 4.63E-19 | 34 |  | 5-fluorouracil | 1.50E-06 | 8 |
| torin1 | 2.73E-16 | 15 |  | MYCN | 5.95E-06 | 7 |
| metribolone | 2.67E-15 | 24 |  | HDAC4 | 2.03E-05 | 5 |
| FMR1 | 6.96E-15 | 16 |  | ammonia | 3.56E-05 | 2 |
| RICTOR | 5.47E-14 | 19 |  | LONP1 | 3.88E-05 | 4 |
| sirolimus | 9.3E-14 | 25 |  | ST1926 | 4.87E-05 | 5 |
| BDNF | 1.28E-11 | 18 |  | TLE3 | 5.46E-05 | 4 |
| **Diseases or Functions Annotation: Epilepsy** | **p-value** | **# Hits** |  | **Diseases or Functions Annotation** | **p-value** | **# Hits** |
| Seizure disorder | 4.07E-14 | 29 |  | Decay of mRNA | 1.30E-08 | 7 |
| Seizures | 4.47E-12 | 25 |  | Synthesis of protein | 2.35E-08 | 13 |
| Infantile or early childhood epileptic encephalopathy | 6.86E-12 | 12 |  | Expression of protein | 1.14E-07 | 11 |
| Developmental epileptic encephalopathy | 7.69E-12 | 12 |  | Parkinson's disease | 1.30E-07 | 9 |
| Familial generalized epilepsy | 4.56E-11 | 12 |  | Nonsense-mediated mRNA decay | 2.66E-07 | 6 |
| Early-onset epilepsy | 9.62E-11 | 13 |  | Progressive encephalopathy | 1.25E-06 | 14 |
| Tonic-clonic seizure | 2.68E-10 | 13 |  | Movement Disorders | 1.26E-06 | 16 |
| Childhood epilepsy | 9.85E-10 | 12 |  | Progressive neuromuscular disease | 1.26E-06 | 10 |
| Epilepsy | 1.34E-09 | 19 |  | Initiation of translation of protein | 1.48E-06 | 6 |
| Epilepsy or neurodevelopmental disorder | 4.79E-09 | 21 |  | Disorder of basal ganglia | 1.64E-06 | 13 |

Altogether, GO and pathway analyses suggest that the deletion of Astro-CaN affects, among others, mitochondrial function, protein synthesis and mTOR signalling, showing significant association with neurological diseases, including AD and epilepsy.

### ACN-KO proteome shows significant overlap with 5xFAD mouse AD model and with human drug-resistant epilepsy datasets

Given the emergence of AD and seizures, we compared all our HIP ACN-KO DEPs with published AD and epilepsy proteome datasets. For AD comparison, we chose a proteome dataset obtained on 8 months-old 5xFAD mouse AD model, which represents the early stage of AD-like pathology (Neuner et al., 2017). HIP ACN-KO dataset showed significant overlap with 5xFAD-vs-NonTg proteome reported by Neuner et al. (18 overlapped proteins, p = 7.02E-15; Fig. 4A, and Supplementary Table 8A). For comparison with epileptic brains, a dataset contributed recently by Karen-Aviram and colleagues (Keren-Aviram et al., 2018) was chosen. This dataset features DEPs from the resected high frequency epileptic foci compared with biopsies of non-epileptic regions from the same patients with drug-resistant epilepsy (denominated as HE). Of 155 DEPs of HIP ACN-KO list, 26 DEPs (16.8%) were common with HE showing significant overlap (p = 1.07E-24; Fig. 4B, Supplementary Table 8B).

**Figure 4.**
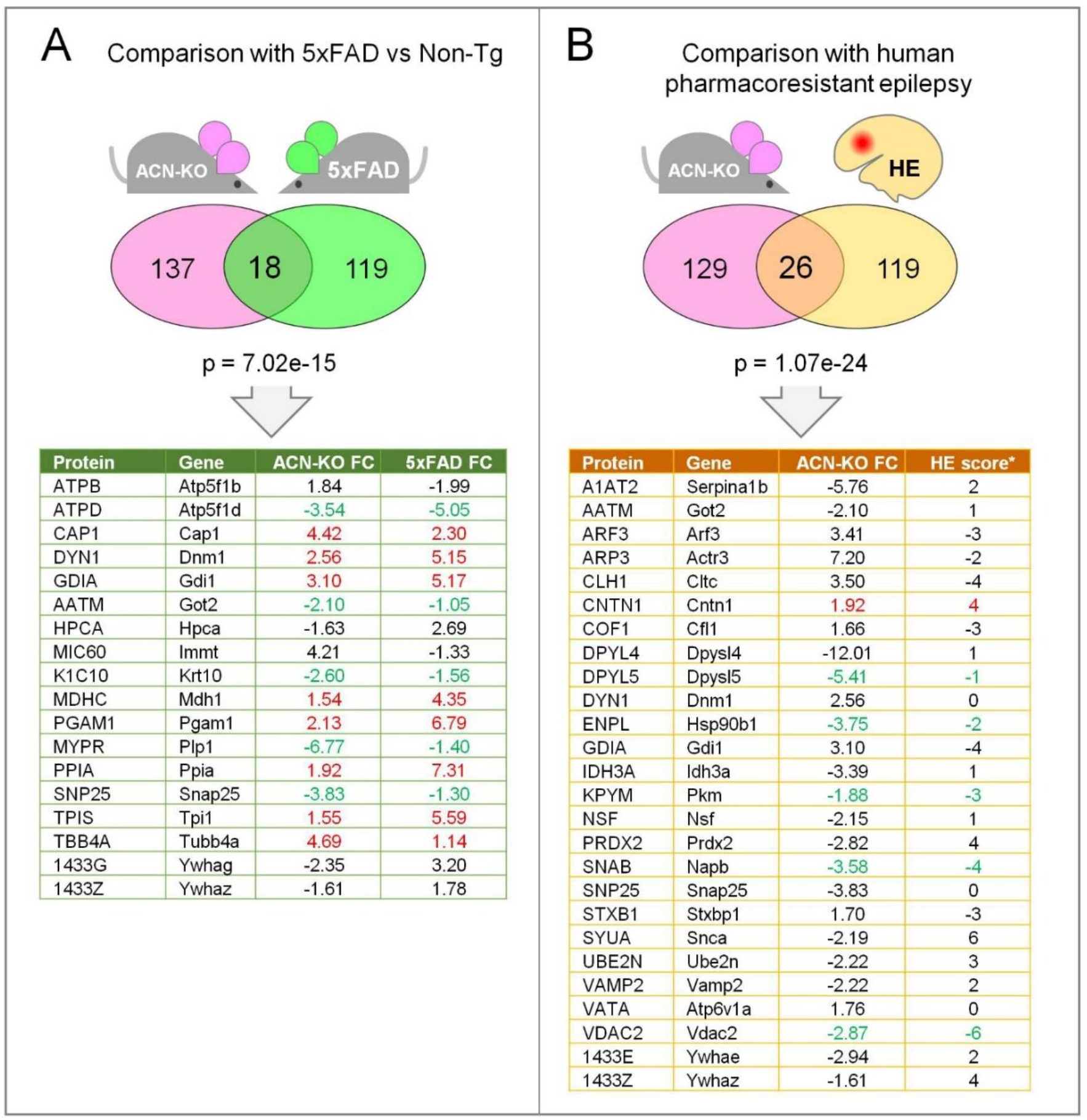
Comparison with 5xFAD mouse and human drug-resistant epilepsy proteome datasets. (A) Comparison with 8 mo old 5xFAD vs NonTg hippocampal proteome (18 common DEPs, listed in the table below; p = 7.02E-15) (Neuner et al., 2017). (B) Comparison with human drug-resistant epilepsy dataset contributed by (Keren-Aviram et al., 2018) obtained on brain biopsies from high-frequency epileptic foci vs low-frequency regions from the same patients. 26 common proteins were found (p = 1.07E-24) listed in the table below. Colour labels: in red co-upregulated; in green: co-downregulated DEPs. Magenta, ACN-KO vs ACN-Ctr DEPs; green, 5xFAD vs NonTg DEPs; light-yellow, human epileptic (HE) DEPs. HE score was calculated as an algebraic sum of numerical values (from 1 to 2 for up-regulated proteins and from −1 to −2 for down-regulated proteins) assigned to DEPs categories according to Keren-Aviram et al., 2018.

Collectively, our omics data suggest that the deletion of CaN from astrocytes, at 1 mo of age, produces deregulation of protein expression resembling that of AD and epilepsy.

### The absence of ultrastructural alterations in ACN-KO hippocampus

Previously, we reported no alterations in neurodevelopment and brain cytoarchitecture of ACN-KO mice (Tapella et al., 2020), while bioinformatic analysis already at 1 month of age ACN-KO mice shows a proteome fingerprint of neurological diseases which may result in alterations at the synaptic level in the hippocampus. Therefore, we have investigated if there were ultrastructural alterations in the hippocampus of ACN-KO mice with consequent behavioural characterization including a hippocampus-dependent task. Transmission electron microscope analysis revealed no ultrastructural alterations in neuropil of CA1 hippocampal region including synapses and mitochondria (Supplementary Figure 8A). Likewise, quantification of dendritic spines in the hippocampus revealed no differences in spine density (p = 0.78, n = 12 for ACN-Ctr, n = 15 for ACN-KO), confirming that, at 1 mo of age, the deletion of CaN from astrocytes does not alter structural and morphological aspects of the CNS (Supplementary Figure 8B).

### ACN-KO mice adopt serial search strategy in Barnes maze

Next, we subjected mice to behavioural tests to assess if general parameters of behaviour and memory of ACN-KO mice were different from that of ACN-Ctr. Results of open field test showed that the distance travelled, the time spent and the average speed of movement in the external and internal parts of the field did not differ between ACN-Ctr and ACN-KO mice (p = 0.73, n = 15) (Suppl. Fig.). Similarly, no changes were found in nesting behaviour (p = 0.145, n = 11), hanging (p = 0.56, n = 8) and tail suspension tests (p = 0.86, n = 18). This indicates that ACN-KO mice do not have deficit of motor activity, equilibrium or muscle strength, neither show anxious or depressive behaviour (Supplementary Figure 9). Next, we subjected ACN-Ctr and ACN-KO mice to Barnes maze behavioural paradigm (Fig. 5A). No differences were observed across acquisition training (AT) sessions in primary latency (F(1,18) = 0.0005, p = 0.98, n = 10 per genotype; Fig. 5), entry latency (F(1,18) = 0.45, p = 0.51) and entry errors (F(1,18) = 0.43, p = 0.52). However, in AT1, ACN-KO mice made significantly more primary errors (13.6 ± 3.05 pokes for ACN-KO vs 6.0 ± 1.03 pokes for ACN-Ctr; p = 0.023) resulting in significative association with genotypes in two-way ANOVA for repeated measures (RM-ANOVA) (F(1,18) = 8.53; p = 0.0091). The difference in the number of primary errors was not due to alteration in locomotor activity as there were no differences in distance travelled (p = 0.57) and average speed (p = 0.45) during AT1 session. These results suggest that, in spite of the difference in the number of primary errors during AT1 session, ACN-KO mice learned the task and succeeded to locate the target hole with the same temporal efficiency as ACN-Ctr mice.

**Figure 5.**
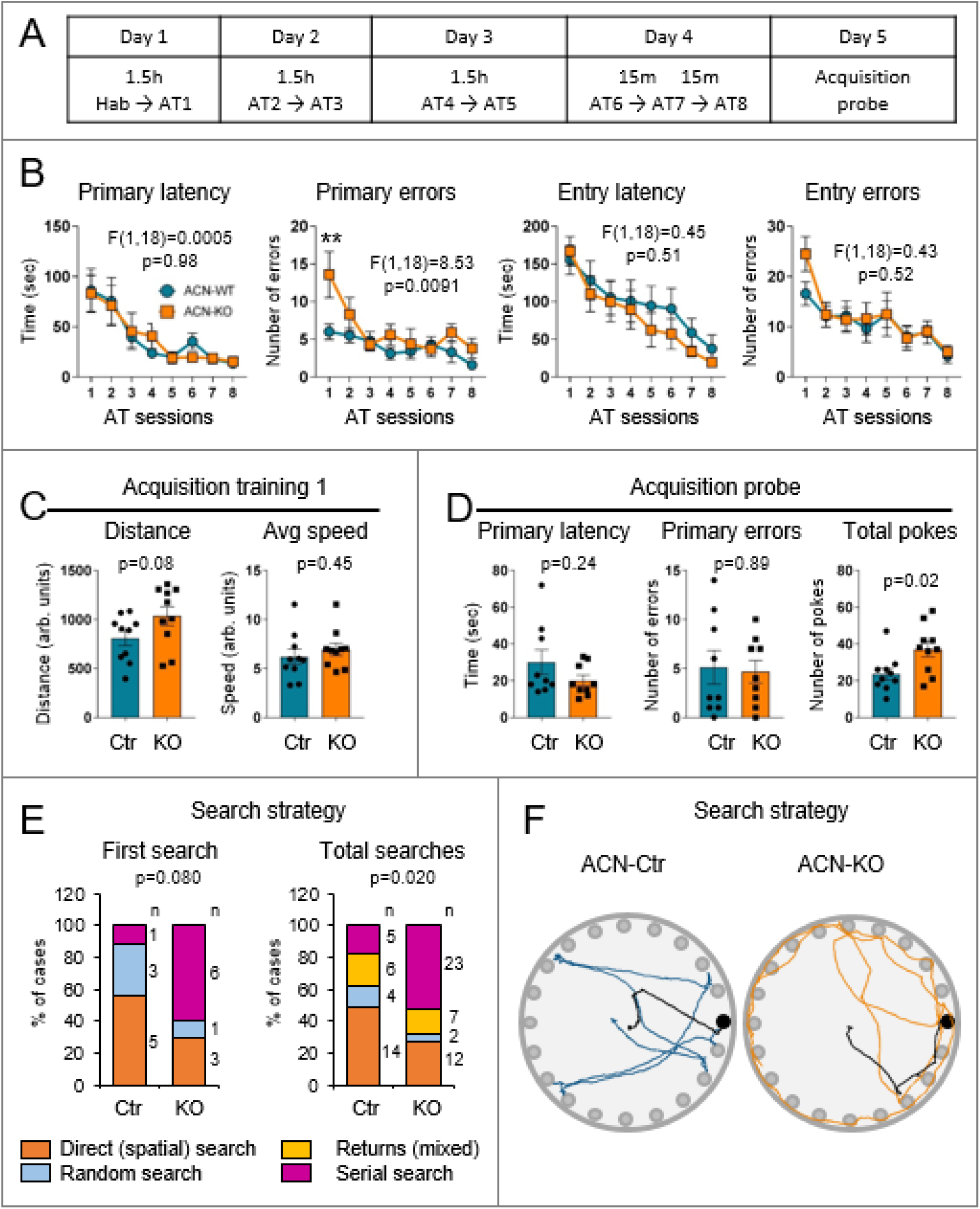
ACN-KO mice learned the Barnes maze task, but use serial search strategy. (A) schematic representation of Barnes maze experiment protocol (Hab = habituation; AT = acquisition training). (B) Performance of ACN-Ctr and ACN-KO mice across AT sessions: no differences were found in primary latency (F(1,18) = 0.0005; p = 0.98), entry latency (F(1,18) = 0.45; p = 0.51) and entry errors (F(1,18) = 0.43; p = 0.52), in primary errors, only in AT1 session, significative difference between ACN-Ctr and ACN-KO mice was found (p = 0.0091). (C) the absence of significative differences in total distance travelled and average (Avg) speed, during AT1 (respectively: p = 0.57; p = 0.45), shows that there are no alterations in general motor and explorative activity in ACN-KO compare to ACN-Ctr mice. (D) ACN-Ctr and ACN-KO mice do not show differences, in the acquisition probe, considering both primary latency (p = 0.19) and primary errors (p = 0.83), but there is a significant difference in the number of total pokes during acquisition probe (p = 0.019). (E) Comparison of search strategies adopted by ACN-Ctr and ACN-KO mice during acquisition training: considering both first search and total searches strategies ACN-Ctr prefer direct search (orange), while ACN-KO prefer serial search (Magenta). (F) representative traces of ACN-Ctr (blue) and ACN-KO (orange) mice; first search is black coloured.

During the acquisition probe (AP) session, in which the escape box was removed and mice were allowed to explore the maze for 90 sec., the primary latency (p = 0.24) and the number of primary errors (p = 0.89) to find the target hole in 90 sec were not different between ACN-Ctr and ACN-KO mice. However, the number of total pokes during the session were significantly higher in ACN-KO than in ACN-Ctr mice (37.1 ± 1.16, n = 10 vs 21.1 ± 3.44 pokes, n = 9, respectively; p = 0.02, unpaired t-test; Fig. 5D-a). This may suggest on the differences in search strategy that mice adopt to locate target hole. To test this we have identified the following search strategies: direct (spatial), serial search and random (mixed) (Gawel et al., 2019). To these three classical search strategies, an additional strategy was discriminated when the mouse returned to the target hole from the first or second adjacent holes (called return) (see Methods section for detailed description of search strategies). In course of the AP session, when the first approach to the target hole was considered, only one (11.1%) of ACN-Ctr mice used serial search, while 3 (33.3%) and 5 (55.6%) mice used, respectively, random and direct search. Instead, 6 (60%) ACN-KO mice used serial search strategy during the first search, while 1 (10%) and 3 (30%) mice used, respectively, random and direct search (p = 0.080, χ^2^, Fisher’s exact test; Fig. 5E-a). The trend of ACN-KO mice to use serial search during the acquisition probe was strengthened when the total number of the target hole searches during 90 sec of probe were counted: ACN-Ctr and ACN-KO mice used, respectively, 5 (17.2%) vs 23 (52.3%) times the serial search strategy, 6 (20.7%) and 7 (15.9%) time the random search, 4 (13.8%) vs 2 (4.5%) the return strategy, and 14 (48.3%) vs 12 (27.3%) spatial search to find the target hole. The differences were significative using χ^2^, Fisher’s exact test (p = 0.020; Fig. 5E-b). These results suggest that ACN-KO mice learn the position of target hole using non-spatial but serial search strategy to locate it. Fig. 5C shows representative traces of ACN-Ctr and ACN-KO mice during AP session, from which is evident that the difference in search strategy is reflected by the time spent and the distance travelled by the mouse in the external part or in the center of the maze. We used these parameters to analyse if the serial search is an intrinsic property of ACN-KO mice linked to anxiety and is independent of learning, or this is the strategy that mice develop during the learning process because they are unable to adopt spatial search strategy. For this, the maze was divided in two zones, external and internal (Fig. 6A), and the distance and time spent in each zone was measured in two occurrences in which all mice spent equal time searching the maze, the habituation session, e. g., before learning, and the AP session. While there was no difference in total distance travelled during both habituation (p = 0.57) and AP session (p = 0.084) (Fig. 6B), ACN-KO mice travelled significantly more (p = 0.0092) (Fig. 6C) and spent more time (p = 0.011) (Fig. 6D) in the external part of the maze than in the internal part in comparison with ACN-Ctr during the AP session but not the habituation.

**Figure 6.**
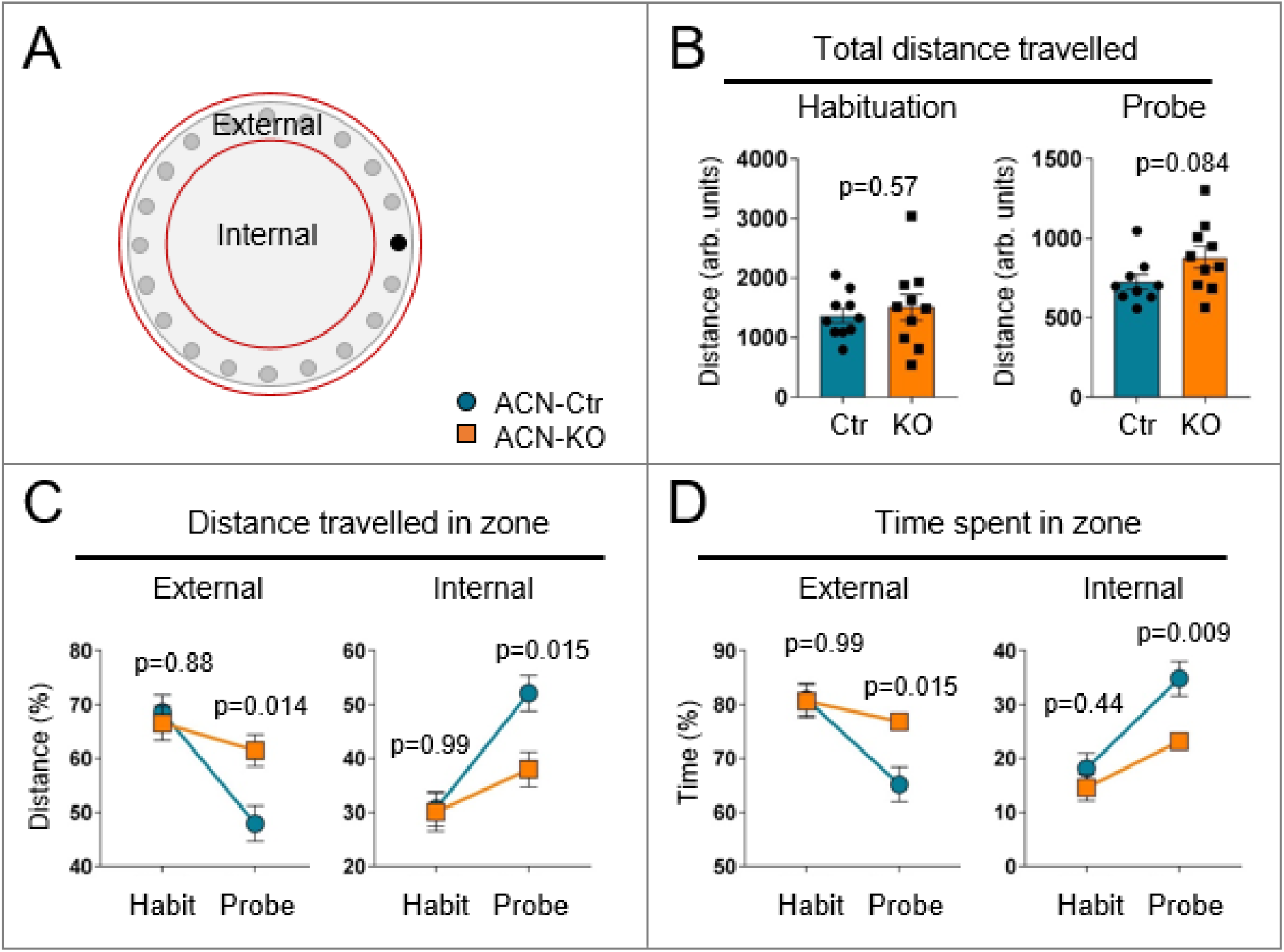
Search strategies alteration is not an intrinsic feature of ACN-KO mice. (A) schematic representation of Barnes maze division in internal zone and external zone. (B) no differences between ACN-Ctr and ACN-KO mice emerged in the total distance travelled during, both, habituation and acquisition probe (respectively: p = 0.57; p = 0.084). (C) % of distance travelled by ACN-Ctr and ACN-KO mice, in external and internal zone, in the habituation (Hab) and probe sessions. (D) % of time spent by ACN-Ctr and ACN-KO mice, in external and internal zone, in the habituation (Hab) and probe sessions.

Taken together, these results show that ACN-KO mice do not have motor alterations or anxiety while presenting subtle alterations in spatial learning.

### ACN-KO mice develop tonic-clonic seizures and neuroinflammation starting from 5 months of age

The emergence of annotations related to epilepsy prompted us to investigate if there was an increased incidence of seizures during life-time of ACN-KO mice. Monitoring of ACN-KO mice revealed that starting from about 5 months of age, both males and females of ACN-KO mice showed increasing risk to develop seizures. Seizures started either upon cage opening, or upon bothering of a mouse by slightly pulling the tail (Supplementary Video 1). The intensity of seizures corresponded to stages from 3 (forelimb clonus) to 5 (rearing and falling with forelimb clonus) according to Racine scale (Racine, 1972) (Supplementary Video 1) or to stages 6-7 according to Pinel and Rovner scale (forelimb clonus with multiple rearing and falling; jumping) (Pinel & Rovner, 1978) (Supplementary Video 2). The percentage of mice developing seizures grew steadily until 10 months after which the incidence did not increase further (Fig. 7A). There were no differences in the risk of seizure development between the three experimental groups, males (n = 50), non-breeding females (n = 36) and breeding females (n = 80) (long-rank Mantel-Cox test, χ^2^ = 0.856, df = 2, p = 0.6518) reaching 40% risk for males and 45% for females (Fig. 7A). We next attempted registration of EEG in ACN-KO males. However, none of mice developed seizure after implantation of electrodes (n = 4) which made it impossible to register EEG during seizures. Thus, we proceeded with registration of seizure-free mice. In all cases registered from ACN-KO mice there was a presence of aberrant electrical activity (100.8 ± 19.25 spikes/24h in ACN-KO vs 3.5 ± 0.50 spikes/24h in ACN-Ctr, p = 0.042, n = 5) resembling that of interictal spikes (Lévesque et al., 2018) (Fig. 7B).

**Figure 7.**
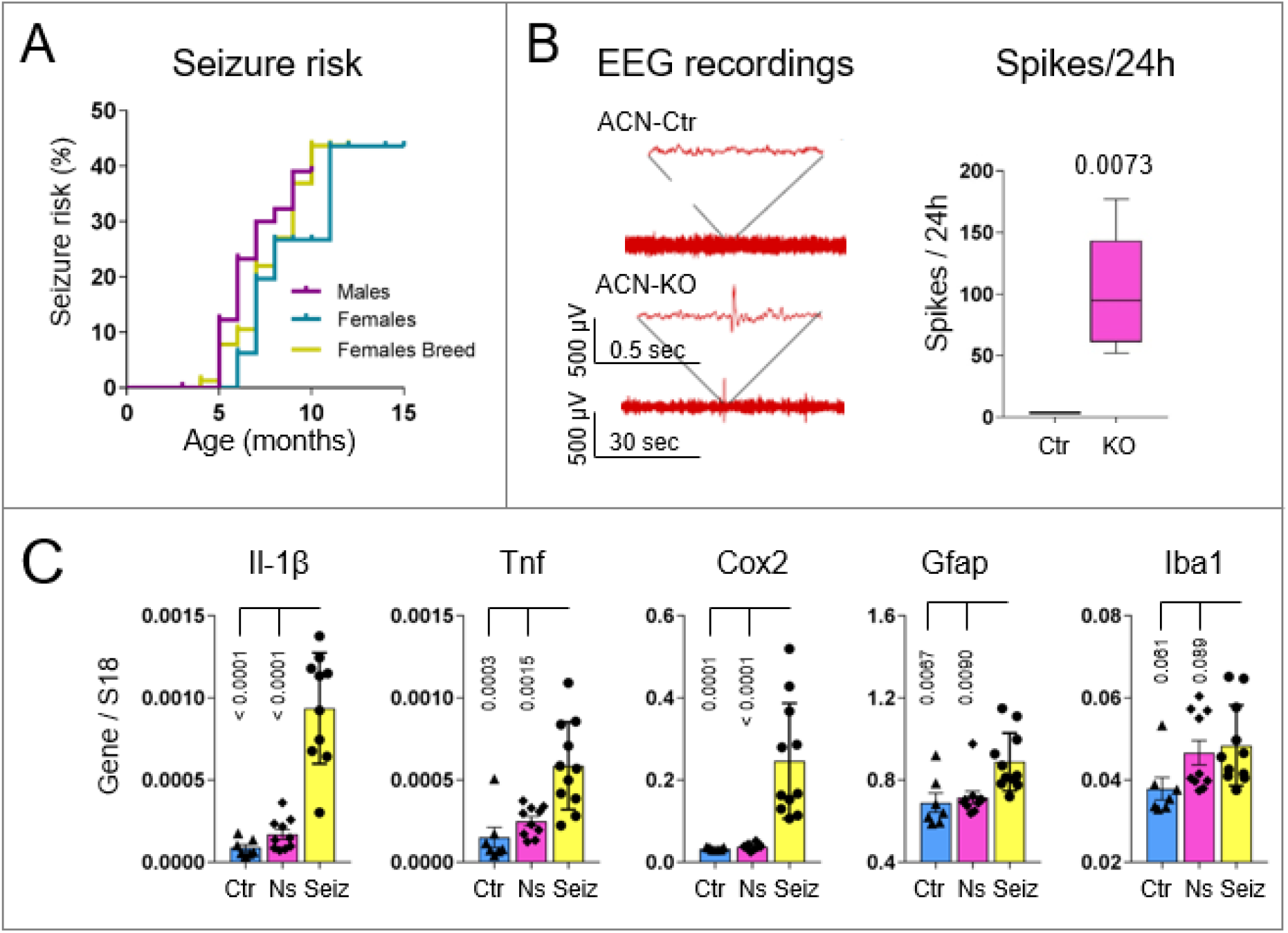
Increased risk to develop tonic-clonic seizures in ACN-KO mice beginning from 5 months of age, and inflammatory phenotype in 9 months-old ACN-KO mice with seizures. (A) Starting from 5 mo of age, both males and females exhibit increasing risk of spontaneous tonic-clonic seizures (violet, males, n = 50; turquoise, females, n = 36; yellow, females in breeding, n = 80, long-rank Mantel-Cox test, χ^2^ = 0.856, df = 2, p = 0.6518). (B) No seizures have been observed in mice which underwent surgery for electrode implantation. However, in seizure-free mice there were aberrant EEG spikes as registered by 24 h recording (100.8 ± 19.25 spikes/24h in ACN-KO vs 3.5 ± 0.50 spikes/24h in ACN-Ctr, p = 0.042, n = 5). (C) Hippocampi were dissected from ACN-Ctr (Ctr, cyan columns), ACN-KO mice which never had seizures (Ns, magenta columns) and from ACN-KO mice with seizures less than 5 min before sacrifice (Seiz, yellow columns). RNA was extracted and real-time PCR was performed using specific primers for pro-inflammatory cytokines (Il-1β, Tnf and Cox2) and markers of gliosis (Gfap and Iba1). Note upregulation of pro-inflammatory and gliotic markers specifically in ACN-KO mice with seizures (p < 0.0001, Seiz vs Ctr and Ns for Il-1β; p = 0.0003 and p = 0.0015, Seiz vs Ctr and Ns, respectively for Tnf; p = 0.0001 and p < 0.0001, Seiz vs Ctr and Ns, respectively for Cox2; p = 0.0067 and p = 0.0090, Seiz vs Ctr and Ns, respectively for Gfap; and p = 0.061 and p = 0.089, Seiz vs Ctr and Ns, respectively for Iba1).

Neuroinflammation is a key feature of human seizure disorders and mouse models of epilepsy (van Vliet et al., 2018). Therefore, we investigated if in ACN-KO mice with seizures this key feature of epileptic brain was respected. qPCR experiments on mRNA extracted from hippocampi of ACN-Ctr (labelled as Ctr in Fig. 7C), ACN-KO mice without seizures (labelled as Ns in Fig. 7C) and ACN-KO mice with recurrent seizures (labelled as Seiz in Fig. 7C) demonstrate that Il-1β, TNF, Cox2 and Gfap mRNA were significantly upregulated in ACN-KO mice with seizures (15.6 fold, p < 0.0001; 3.9 fold, p = 0.0003; 7.62 fold, p = 0.0001 and 1.29 fold, p = 0.0067, compared with ACN-Ctr, respectively), indicating an ongoing neuroinflammatory process. Together, our data suggest that the deletion of CaN from astrocytes results in development of delayed spontaneous tonic-clonic seizures from 5 months of age.

## DISCUSSION

The present study was designed to investigate the effect(s) of the deletion of Astro-CaN on global transcriptional and translational remodelling in the CNS of ACN-KO mice at 1 month of age, a timepoint when the CaN KO is fully expressed, development and neuronal differentiation in the CNS are completed, while there are minimal compensatory effects of the deletion of Astro-CaN. The principal findings of this work are: (i) in resting conditions, Astro-CaN does not play a significant role in transcription; (ii) at 1 month of age, protein expression was altered in hippocampus and cerebellum of ACN-KO mice in a brain area-specific manner. In addition, a different set of proteins was found altered when specifically looking at hippocampal and cerebellar synaptosomes; (iii) changes in protein expression occurred in all principal cellular types of the CNS including astrocytes, neurons, oligodendrocytes, microglia and endothelial cells; (iv) bioinformatic analysis revealed overrepresentation of differentially expressed proteins related to CNS disorders, specifically to AD and seizure disorder; (v) comparisons with published proteome datasets indicate significant overlap of HIP ACN-KO dataset with those related to AD and epilepsy; (vi) ACN-KO mice do not exhibit alterations in locomotor activity, signs of anxiety of depression; (vii) in Barnes maze, ACN-KO mice were able to locate the target hole with the same efficiency as ACN-Ctr mice but used non-spatial search strategy, serial search; and (vii) beginning from about 5 months of age ACN-KO mice show increased risk to develop tonic-clonic seizures.

### Posttranscriptional nature of the Astro-CaN deletion effects

In astrocytes, CaN activity is mainly associated with transcriptional remodelling through activation of transcription factors NFAT and NF-kB. Such a transcriptional remodelling has been shown in several pathological conditions *in vivo* and *in vitro* in response to toxic or pro-inflammatory stimuli (Fernandez et al., 2012; Furman & Norris, 2014; Lim, Rocchio, et al., 2016). Neuronal activity-dependent activation of CaN followed by NFAT nuclear translocation has been demonstrated in pericytes after field stimulation of hippocampal slices (Filosa et al., 2007), while *in vitro* our group has shown a robust Astro-CaN activation in mixed neuron-astrocytes hippocampal cultures upon chemical induction of long-term potentiation (LTP) (Lim et al., 2018). Here we show that the remodelling of CNS proteome, as a result of the deletion of CaN from astrocytes, occurs in the absence of transcriptional alterations, suggesting that the principal mean by which CaN acts in healthy astrocytes is through protein de-phosphorylation. However, it should be emphasized that for RNA-Seq and SG-MS analyses we used resting, not stimulated and not stressed animals. Therefore, at least in principle, a stress-or-stimulation-induced Astro-CaN-mediated transcriptional activity, as it occurs in neurons during late phase of LTP (Baumgärtel & Mansuy, 2012), cannot be *a priori* ruled-out. Nevertheless, we can speculate that, upon continuous modifications in homeostatic demand during dynamic neuronal activity, fast dephosphorylation/phosphorylation signalling would be more advantageous compared to a relatively slower process such as transcription.

### Astro-CaN deletion predicts neuropathology, does it mean that neuropathology is caused by Astro-CaN dysfunction?

The central finding of this work is that, in response to the deletion of CaN from astrocytes, the molecular signature of neuropathology was evident already at 1 month of ACN-KO mouse age. As shown in the gene ontology analysis, the molecular signature resembles that of several neurological and neurodegenerative diseases, e.g., OXPHOS alterations were common between PD, HD and AD.

However, IPA pathway analysis indicates on a particular link with AD. Strikingly, the top 3 Upstream Regulators found by IPA analysis were three the major causes of familiar AD, namely Tau (MAPT), PS1 and APP with 47 common proteins downstream of these hypothetical regulators (30.3% of HIP list, Supplementary Table 7B). Because of FAD mutated proteins have been traditionally regarded as neuronal proteins, one could draw a straightforward conclusion that the deletion of CaN from astrocytes predisposes neurons to harmful effects of FAD mutations partly recreating the fingerprint of FAD-related proteome. Analysis of cell-specificity of this list shows that the majority of proteins are, indeed, neuron-specific or enriched in neurons (Fig. 3B).

Of special interest is the overlap with a mouse model of incipient AD, 8 months-old 5xFAD mice (Neuner et al., 2017). However, since ACN-KO mice do not bear FAD mutations, it is unlikely that ACN-KO mouse represents an AD mouse model. Instead, significant overlap with hippocampal proteome of 5xFAD mice suggest that the alterations of Astro-CaN activity may occur during AD pathogenesis. Moreover, we propose that it may constitute a common denominator for a group of neuropathological diseases as suggested by bioinformatic analysis. In this light, considering the increased risk of seizures in AD patients (Nicastro et al., 2016; Vossel et al., 2017), the association of Astro-CaN KO specifically with AD and epilepsy assumes particular significance. Therefore, investigation of Astro-CaN activity and its association with central and peripheral markers at early disease stages may lead to the development of novel approaches in diagnosis, prevention and therapy.

### Subtle alterations in spatial memory in ACN-KO mice

Interestingly, at 1 mo of age, we did not find ultrastructural alteration in the hippocampal neuropil of ACN-KO mice, neither they have alterations in basic behavioural performances like locomotor activity, equilibrium and strength. In addition, ACN-KO mice did not exhibit anxious o depressive behaviour, as none of the parameters in open field, nesting or tail suspension tests was changed. However, alterations in navigational strategies which ACN-KO mice adopt to locate the target hole in Barnes maze test suggest a functional impairment of hippocampal circuits. Barnes maze is considered to be a hippocampus-dependent task in which the ability to use spatial memory is tested (Sharma et al., 2010). However, a mouse is able to find the target hole even without spatial orienteers, using the information inherent to maze exploring holes serially (Harrison et al., 2006). In this terms, one could conclude that ACN-KO mice, which adopt serial search to locate the target hole, have clear functional hippocampal impairment. However, lately, it has been argued that hippocampus is used for non-spatial memory processing due to its ability to time-parse elementary memory events to integrate temporal (when), episodic (what), spatial (where) information into memory traces (Eichenbaum, 2017; Sugar & Moser, 2019). In light of this paradigm it is plausibly to conclude that ACN-KO mice fail to develop spatial search and avail to serial exploration of holes due to functional dissociation of temporal and spatial representations both of which are hippocampus-dependent (Eichenbaum, 2017). Such a dissociation has been found in the hippocampus of an AD mouse model, rTg4510 mice, in which temporal sequences of neuronal firing can persist while spatial firing is disrupted (Cheng & Ji, 2013). This rational is supported by several studies of animal models of AD and epilepsy. Thus, Macedo and colleagues (Macêdo et al., 2018) showed that rats, after hippocampal infusion of β-amyloid peptides, used non-spatial strategies to solve the maze; similarly, in kainic acid model of temporal lobe epilepsy, mice understood the context and solved the maze using serial but not spatial search (Van Den Herrewegen et al., 2019), suggesting that the inability to form spatial representations may be a common feature between AD and epilepsy.

Striking similarity was found in Barnes maze performance, specifically in the acquisition training profiles and the preference of serial search strategy, between ACN-KO mice and a mouse with neuronal expression of Ca^2+^-insensitive mutant of Ca^2+^/calmodulin-dependent kinase II (CaMKII) in which long-term potentiation in theta-range (5-10 Hz) of stimulating frequencies was specifically impaired (Bach et al., 1995). Speculatively, this may suggest a mechanistic link between Astro-CaN-regulated CNS proteostasis and neuronal activity and plasticity. Further thorough molecular and electrophysiological analysis of neuronal activity and plasticity is needed to investigate the link between Astro-CaN and neuronal CaMKII. While such investigation is beyond the aim of the present contribution, the link between neuronal excitability, found to be impaired in ACN-KO mice (Tapella et al., 2020), and the activity of CaMKII in neurons is well documented (Liu & Murray, 2012).

### Is ACN-KO mouse a novel model of spontaneous seizures and epilepsy?

Another important finding of this contribution regards the increased risk of ACN-KO mice to develop motor seizures in the period from 5 to 12 months of age. As we have previously reported, to 20^th^ postnatal day ACN-KO mice develop full CaN KO in the hippocampus and the cerebellum (Tapella et al., 2020). Therefore, it is reasonable to suggest that the seizures are unlikely to be the direct effect of Astro-CaN KO. A delay of 5-10 month from the CaN-KO till seizure development suggests that a constellation of long-lasting alterations, downstream of Astro-CaN, may lead to seizure development. While the primary cause is represented by the Astro-CaN KO, the very cause of seizures is currently unknown. At the time of seizures ACN-KO mice show inflammatory phenotype closely resembling that observed in human disease and animal models of epilepsy (van Vliet et al., 2018). However, despite that the onset and development of epilepsy are strongly associated with neuroinflammation (Vezzani et al., 2019), it is prematurely to conclude that the seizures in ACN-KO mice are caused by the inflammatory process. Moreover, at 1 month of age of ACN-KO mice, the molecular epileptic fingerprint does not evidence inflammatory component but strongly suggests the insufficiency of glial support to neurons because: (i) primary cause is the deletion of an astroglial enzyme; (ii) altered expression of astrocyte-specific membrane transporters as Glast (Dematteis et al., 2020), Glt-1 and Atp1a2; and (iii) overrepresentation of DEPs related to myelin sheath suggesting inadequate oligodendrocytic function. In support of this, compromised myelinisation of axons in hippocampus has been repeatedly associated with insurgence of seizure (Song et al., 2018; Ye et al., 2013). In this light, the significant overlap 1 mo-old ACN-KO mice proteome with that from human patients with intractable epilepsy (Keren-Aviram et al., 2018) is of special interest. Indeed, Keren-Aviram and colleagues reported downregulation of GFAP in the epileptic foci and did not report appearance of other makers of neuroinflammation.

Lastly, a “spontaneous” nature of seizures discriminates ACN-KO mice from the majority of animal models of epilepsy (Kandratavicius et al., 2014). We found that two third of ACN-KO mice do not develop seizures during their life-time, and the appearance of seizures cannot be predicted with absolute certainty. Manipulation with ACN-KO mice and/or surgical procedures decreases the probability to develop seizure to the extent that we were not able to register seizures in the electrode-implanted animals. All this renders ACN-KO mice time and cost-ineffective as a model of epilepsy (Kandratavicius et al., 2014). However, “fast” and cost-effective animal models are not able to faithfully reproduce all aspects of the human seizure disorder (Kandratavicius et al., 2014). This is particularly true for drug-resistant forms of epilepsy (Tang et al., 2017). We, therefore, envisage that the results obtained from the thorough characterisation of epileptic phenotype of ACN-KO mice may outperform the time-and cost-related disadvantages of this novel and spontaneous model of seizure disorder.

### Conclusion: Is Astro-CaN a master-regulator of homeostasis in the CNS?

The observations that glial cells change very early during neuropathology (Rodríguez et al., 2014; Verkhratsky et al., 2016) led to hypothesize that an insufficient homeostatic support to other cells in the CNS may be among the primary events which predispose neurons to dysfunction (Verkhratsky et al., 2016, 2019). In this context, we have proposed that deregulation of astroglial Ca^2+^ signalling, downstream of CaN, may contribute to this homeostatic failure (Lim et al., 2014; Lim, Rodríguez-Arellano, et al., 2016). Our present data provide further support to this hypothesis suggesting that Astro-CaN may work as a Ca^2+^-sensitive hub orchestrating homeostatic activities of astrocytes. While molecular and signalling details of CaN activation and of Astro-CaN downstream signalling are a matter of future research, our results provide a framework for investigations suggesting that protein synthesis (Dematteis et al., 2020), myelination and mitochondria, alongside with mTOR signalling and with control of ion homeostasis (Tapella et al., 2020), may represent principal components of Astro-CaN-controlled homeostatic network.

## METHODS

### Generation and handling of ACN-KO mice

Generation and handling of conditional CaN knockout (KO) in GFAP-expressing astrocytes (<u>a</u>stroglial <u>c</u>alci<u>n</u>eurin <u>KO</u>, ACN-KO) has been described previously (Tapella et al., 2020). Briefly, CaNB1^flox/flox^ mice (Neilson et al., 2004) (Jackson Laboratory strain B6;129S-Ppp3r1tm2Grc/J, stock number 017692) were crossed with GFAP-Cre mice (Gregorian et al., 2009) (Jackson Laboratory strain B6.Cg-Tg(Gfap-cre)77.6Mvs/2J, stock number 024098). The line was maintained by crossing CaNB1^flox/flox^/GfapCre^-/-^ (thereafter referred as to ACN-Ctr) males with CaNB1^flox/flox^/GfapCre^+/-^ (ACN-KO) females.

Mice were housed in the animal facility of the Università del Piemonte Orientale, with unlimited access to water and food. Animals were managed in accordance with European directive 2010/63/UE and with Italian law D.l. 26/2014. The procedures were approved by the local animal-health and ethical committee (Università del Piemonte Orientale) and were authorized by the national authority (Istituto Superiore di Sanità; authorization numbers N. 77-2017 and N. 214-2019). All efforts were made to reduce the number of animals bred and used.

### Next Generation RNA sequencing (RNA-Seq)

For RNA-Seq, total RNA from hippocampal and cerebellar tissues were extracted using the SPLIT RNA Extraction Kit (Cat. 00848; Lexogen GmbH, Vienna, Austria) according to the manufacturer’s instructions. 200 ng of RNA were subject to an Illumina sequencing using Lexogen QuantSeq FWD - SR75 - DA Service. Reads were mapped to the reference genome using STAR aligner v2.7 (Dobin et al., 2013), while the differential expression analysis was performed with DESeq2 v1.26 Bioconductor package (Love et al., 2014). Six (6) animals per condition were independently processed from 1-month old ACN-Ctr and ACN-KO mice.

### Isolation of total RNA and real-time PCR

For validation of RNA-Seq and shotgun proteomic results, samples were prepared from a separate set of 12 ACN-Ctr and ACN-KO 1 month old mice.

For real-time PCR, hippocampal or cerebellar tissues were lysed in Trizol reagent (Invitrogen, Cat. 15596026). Total RNA was extracted using according to manufacturer’s instructions using chloroform extraction followed by isopropanol precipitation. After washing with 75% ethanol, RNA was dried and resuspended in 20-30 μL of RNAse-free water. 0.5-1 μg of total RNA was retrotranscribed using SensiFAST cDNA Synthesis Kit (BioLine, London, UK). Real-time PCR was performed using iTaq qPCR master mix according to manufacturer’s instructions (Bio-Rad, Segrate, Italy, Cat. 1725124) on a SFX96 Real-time system (Bio-Rad). To normalize raw real-time PCR data, S18 ribosomal subunit was used. Sequences of oligonucleotide primers is listed in Supplementary Methods Table 1. The real-time PCR data are expressed as delta-C(t) of gene of interest to S18 allowing appreciation of the expression level of a single gene.

### Preparations of tissue homogenates and synaptosomal fractions

Tissues were homogenized using glass-Teflon 1 ml homogenizer in lysis buffer (50 mM Tris HCl (pH 7.4), sodium dodecyl sulfate (SDS) 0.5%, 5 mM EDTA). Synaptosomal fractions were prepared as reported previously (Gillardon, 2006) from hippocampal or cerebellar tissues pooled from two 1 month old ACN-Ctr or ACN-KO mice. Briefly, mice were anesthetized by intraperitoneal injection of Zoletil (80 mg/kg) and Xylazine (45 mg/kg) and sacrificed by decapitation. Hippocampi and cerebella were rapidly dissected and placed into ice-cold homogenization buffer containing 50 mM MOPS, pH 7.4, 320 mM sucrose, 0.2 mM DTT, 100 mM KCl, 0.5 mM MgCl2, 0.01 mM EDTA, 1 mM EGTA, protease inhibitor cocktails (PIC, Calbiochem) and phosphatase inhibitor Na3VO4 (1μM). All subsequent steps were performed at 4°C. The tissues were minced and homogenized in 1:10 w/v homogenization buffer with 12 strokes in a Teflon-glass douncer. The homogenates were then centrifuged for 10 min at 800g followed by centrifugation of the supernatant at 9200g for 15 min. The resulting P2 pellet, representing the crude synaptosomal fraction, was solubilized in lysis buffer.

### Western blot

20 μg proteins were resolved on 8% of 12% SDS-polyacrylamide gel. Proteins were transferred onto nitrocellulose membrane using Trans-Blot Turbo Transfer System (Bio-Rad), blocked in skim milk (5%, for 1 h; Cat. 70166; Sigma) and immunoblotted with indicated antibody. Mouse anti-Tubulin (1:5000, Cat. T7451) or anti-β-actin (1:10000, Cat. A1978), both from Sigma (Milan, Italy) were used to normalize protein load. Primary antibodies were: mouse anti-Sy38 (1:500, Cat. MAB-10321); rabbit anty-Cox6a1 (1:500, Cat. AB-83898); rabbit anti-Vdac2 (1:500, Cat. AB-83899); rabbit anti-Atp5j (1:500, Cat. AB-83897); rabbit anti-Dpysl4 (1:500, Cat. AB-83901); rabbit anti-Gfap (1:500, Cat. N. AB-10682), all from Immunological Sciences (Rome, Italy); rabbit anty-Rps10 (1:500, Cat. PA5-96460, Thermofisher, Milan, Italy); rabbit anty-Glast (1:500, Cat. NB100-1869, NovusBio, Milan, Italy); rabbit anty-Atp1a2 (1:1000, Cat. ANP-002, Alomone Labs, Jerusalem, Israel); mouse anti-PSD95, clone K28/43 (1: 1000, Cat. MABN68, Merck Millipore, Milan, Italy); mouse anti-Snap25 (1:600, Cat. sc-376713, SantaCruz, Dallas, Texas, USA). Goat anti-mouse IgG (H+L) horseradish peroxidase-conjugated secondary antibody (1: 8000; Cat. 170-6516, Bio-Rad, Milan, Italy) was used. The protein bands were developed using SuperSignal West Pico chemiluminescent substrate (Cat. 34078; Thermofisher). Densitometry analysis was performed using the Quantity One software (Bio-Rad).

### Proteomic analysis

#### In solution digestion

Lysates were digested using the following protocol: samples were prepared to have 100 μg of protein in a final volume of 25 μL of 100 mM NH_4_HCO_3_. Proteins were reduced using 2.5 μL of dithiothreitol (200 mM DTT stock solution) (Sigma) at 90 °C for 20 min, and alkylated with 10 μl of Cysteine Blocking Reagent (Iodoacetamide, IAM, 200 mM Sigma) for 1 h at room temperature in the dark. DTT stock solution was then added to destroy the excess of IAM. After dilution with 300 μL of water and 100 μL of NH4HCO3 to raise pH 7.5-8.0, 5 μg of trypsin (Promega, Sequence Grade) was added and digestion was performed overnight at 37 °C. Trypsin activity was stopped by adding 2 μL of neat formic acid and samples were dried by Speed Vacuum (Rocchio et al., 2019).

The peptide digests were desalted on the Discovery^®^ DSC-18 solid phase extraction (SPE) 96-well Plate (25 mg/well) (Sigma-Aldrich Inc., St. Louis, MO, USA). The SPE plate was preconditioned with 1 mL of acetonitrile and 2 mL of water. After the sample loading, the SPE was washed with 1 mL of water. The adsorbed proteins were eluted with 800 μL of acetonitrile:water (80:20). After desalting, samples were vacuum evaporated and reconstituted with 20 μL of 0.05% formic acid in water. Two μL of stable-isotope-labeled peptide standard (DPEVRPTSAVAA, Val-13C5 15N1 at V10, Cellmano Biotech Limited, Anhui, China) was spiked into the samples before the LC-MS/MS analysis and used for instrument quality control.

#### Label-free proteomic analysis

LC-MS/MS analyses were performed using a micro-LC Eksigent Technologies (Dublin, USA) system with a stationary phase of a Halo Fused C18 column (0.5 × 100 mm, 2.7 μm; Eksigent Technologies, Dublin, USA). The injection volume was 4.0 μL and the oven temperature was set at 40 °C. The mobile phase was a mixture of 0.1% (v/v) formic acid in water (A) and 0.1% (v/v) formic acid in acetonitrile (B), eluting at a flow-rate of 15.0 μL min^-1^ at an increasing concentration of solvent B from 2% to 40% in 30 min. The LC system was interfaced with a 5600+ TripleTOF system (AB Sciex, Concord, Canada) equipped with a DuoSpray Ion Source and CDS (Calibrant Delivery System). Samples used to generate the SWATH-MS (Sequential window acquisition of all theoretical mass spectra) spectral library were subjected to the traditional data-dependent acquisition (DDA): the mass spectrometer analysis was performed using a mass range of 100–1500 Da (TOF scan with an accumulation time of 0.25 s), followed by a MS/MS product ion scan from 200 to 1250 Da (accumulation time of 5.0 ms) with the abundance threshold set at 30 cps (35 candidate ions can be monitored during every cycle). Samples were then subjected to cyclic data independent analysis (DIA) of the mass spectra, using a 25-Da window. A 50-ms survey scan (TOF-MS) was performed, followed by MS/MS experiments on all precursors. These MS/MS experiments were performed in a cyclic manner using an accumulation time of 40 ms per 25-Da swath (36 swaths in total) for a total cycle time of 1.5408 s. The ions were fragmented for each MS/MS experiment in the collision cell using the rolling collision energy. The MS data were acquired with Analyst TF 1.7 (SCIEX, Concord, Canada). Three instrumental replicates for each sample were subjected to the DIA analysis (Dalla Pozza et al., 2018; Martinotti et al., 2016).

#### Protein Database Search

The mass spectrometry files were searched using Protein Pilot (AB SCIEX, Concord, Canada) and Mascot (Matrix Science Inc., Boston, USA). Samples were input in the Protein Pilot software v4.2 (AB SCIEX, Concord, Canada), which employs the Paragon algorithm, with the following parameters: cysteine alkylation, digestion by trypsin, no special factors and False Discovery Rate at 1%. The UniProt Swiss-Prot reviewed database containing mouse proteins (version 12/10/2018, containing 25137 sequence entries). The Mascot search was performed on Mascot v2.4, the digestion enzyme selected was trypsin, with 2 missed cleavages and a search tolerance of 50 ppm was specified for the peptide mass tolerance, and 0.1 Da for the MS/MS tolerance. The charges of the peptides to search for were set to 2 +, 3 + and 4 +, and the search was set on monoisotopic mass. The instrument was set to ESI-QUAD-TOF and the following modifications were specified for the search: carbamidomethyl cysteines as fixed modification and oxidized methionine as variable modification.

#### Protein quantification

The quantification was performed by integrating the extracted ion chromatogram of all the unique ions for a given peptide. The quantification was carried out with PeakView 2.0 and MarkerView 1.2. (Sciex, Concord, ON, Canada). Six peptides per protein and six transitions per peptide were extracted from the SWATH files. Shared peptides were excluded as well as peptides with modifications. Peptides with FDR lower than 1.0% were exported in MarkerView for the t-test.

### Cell specificity analysis

To assess cell specificity, two databases were primed: Human Protein Atlas (HPA, https://www.proteinatlas.org/) and the mouse brain cell-specific RNA-Seq database contributed by (Zhang et al., 2014). HPA immunoreactivity scores in glial cells and neurons were labelled as: NA (not assayed), ND (not detected, score = 0), Low (score = 1), Medium (score = 3) and High (score = 4). Hits with a score = 0 in glial cells and score > 0 in neurons was assigned to as Neuron-specific; hits in which score ≠ 0 and neuron (N)/glia (G) ratio > 1 were assigned to as Enriched in neurons; hits in which score ≠ 0 and N/G ratio = 1 were assigned to as Ubiquitous; hits in which score ≠ 0 and N/G ratio < 1 were assigned to as Enriched in glia; and hits with a score = 0 in neurons and score > 0 in glia were assigned to as Glia-specific.

To score the Zhang et al database, excel files with Mean FPKM cell-specific values were downloaded. For each gene, ratios were calculated as (Neurons)/(All other cell types); (Astrocytes)/(All other cell types); (Oligodendrocyte Precursor Cells + Non-myelinating Oligodendrocytes + Myelinating Oligodendrocytes)/(All other cell types); (Microglia)/(All other cells types). Genes with ratios > 5 were assigned as cell-type specific, and the presence of these genes in HIP ACN-KO list was specified.

### Bioinformatic analysis

For bioinformatic analysis, DEPs from whole tissue preparations were pooled with correspondent synaptosomal DEPs: Wht-Hip U Syn-Hip = HIP; Wht-Cb U Syn-Cb = CB. Wht-Hip and Syn-Hip did not contain common DEPs at |FC| > 1.5 and p-value < 0.05 and were pooled without modifications. Wht-Cb and Syn-Cb (|FC| > 1.5 and p-value < 0.05) contained 3 common DEPs. In this case the DEP with lower p-value regardless of its FC was taken for further analysis. This resulted 155 and 54 unique DEPs in HIP and CB lists, respectively.

#### DAVID gene ontology (GO) analysis

Gene ontology (GO) analysis was performed using The <u>D</u>atabase for <u>A</u>nnotation, <u>V</u>isualization and <u>I</u>ntegrated <u>D</u>iscovery (DAVID) v.6.8 tool (https://david.ncifcrf.gov/) (Huang et al., 2009). Overrepresented GO terms which passed Benjamini correction (p < 0.05) were considered significant.

#### STRING protein-protein interaction analysis

For prediction of protein-protein interactions and clustering using K-means algorithm, <u>S</u>earch <u>T</u>ool for the <u>R</u>etrieval of <u>I</u>nteracting <u>G</u>enes/Proteins (STRING) v10.5 online software was used (https://string-db.org/) (Szklarczyk et al., 2017).

#### IPA pathway analysis

For pathway analysis, Ingenuity Pathway Analysis package (Content version: 49932394; Release Date: 2019-11-14) was used. Ingenuity Canonical Pathways, Disease and Bio Functions, Upstream Regulators and Causal Network Regulators were downloaded using default settings.

### Comparison with AD and epilepsy datasets

HIP ACN-KO differential protein expression was compared with published mouse AD and human epilepsy proteome datasets. The proteomics analysis by Neuner et al. (Neuner et al., 2017) on 8-month-old 5xFAD hippocampus mouse model was used as AD reference and DEPs were matched by primary accession number as retrieved from the UniProtKB/Swiss-Prot reviewed database (release 2020_01). The dataset provided by Keren-Aviram et al. (Keren-Aviram et al., 2018) on human refractory epilepsy was used as epileptic reference. In this case, to allow interspecies comparison, BioMart tool (www.ensembl.org) was used to extract genes that were orthologous between human and mouse. Afterwards, DEPs were again matched by primary accession number. In both the cases, the significance of the overlap between sets was evaluated through hypergeometric test, using as background reference the number of protein-coding transcripts detected by RNA-Seq (N=14,333 genes). Human epilepsy (HE) score shown in Figure 4 and Supplementary Table 8B was calculated as an algebraic sum of numerical values (from 1 to 2 for up-regulated proteins and from −1 to −2 for down-regulated proteins) assigned to DEPs categories according to Keren-Aviram et al., 2018.

### Transmission electron microscopy

ACN-Ctr and ACN-KO mice were deeply anesthetized and perfused through the ascending aorta with phosphate buffered saline (PBS, 0.1 M; pH 7.4) and heparin (1.25 unit/ml) for not more than 2 minutes, followed by a 20-minute perfusion with 2% paraformaldehyde (PFA) and 2.5% glutaraldehyde in PBS. Hippocampi was excised and cut in the sagittal plane with a razor blade and fixed in 4% paraformaldehyde (PFA) and 2% glutaraldehyde in PBS and then for 2 h in OsO_4_. After dehydration in graded series of ethanol, tissue samples were cleared in propylene oxide, embedded in Epoxy medium (Epon 812 Fluka) and polymerized at 60 °C for 72 h. From each sample, one semithin (1 μm) section was cut with a Leica EM UC6 ultramicrotome and mounted on glass slides for light microscopic inspection. Ultrathin (70 nm thick) sections of areas of interest were obtained, counterstained with uranyl acetate and lead citrate, and examined with an Energy Filter Transmission Electron Microscope (EFTEM, ZEISS LIBRA^®^ 120) equipped with a YAG scintillator slow scan CCD camera.

### Golgi stain and dendritic spine density determination

One-month-old male mice were perfused transcardially with 0.9% NaCl in distilled water. Brains were taken and immersed in Golgi-Cox solution of FD Rapid Golgi Stain Kit (Cat. PK 401, FD Neuro Technologies, Columbia, MD) for 1 week, in the dark, at room temperature. After the period of impregnation, brains were immersed in solution C (Cat. PK 401, FD Neuro Technologies, Columbia, MD) and processed according manufacturer’s instructions. The brains were sliced, using a vibratome, at 100 μm thickness; during the operation, the sample was kept wet with solution C (Cat. PK 401, FD Neuro Technologies, Columbia, MD). Then slices were put on gelatinized slides (2% gelatin (Merk), 1% KCr(SO_4_)_2_·12H_2_O (Carlo Erba) and stained in the dark. After washing twice (4 min) with double distilled water, slices were stained in staining solution (10 min), prepared according manufacturer’s instructions. After three double distilled water washes, slides were dehydrated for 4 min in increasing percentages of ethanol (50, 75, and 95%; all Sigma-Aldrich) and washed four times (4 min) in 100% ethanol. Finally, slides were immersed three times (4 min) in xylene and mounted on coverslips with Entellam (Electron Microscopy Sciences). Images were taken with a 63x objective in a white field. Neurons were analyzed with Neuron Studio software and statistical analysis was done in GraphPad Prism v.7.

### Behavioral tests

#### Open field test

Open field test, a measure of exploratory behavior, general activity and anxiety (Gould et al., 2009), was performed using 60 cm (length) x 40 cm (width) x 35 cm (height) boxes made from white high density and non-porous plastic. Mouse was placed in the center of the box and allowed to freely explore the environment for 10 minutes. Mouse movements were registered and processed offline using PC-based software. The area of the box was divided in external and central areas. The following parameters were calculated: total distance travelled (DT), distance travelled in the external part (DE) and in the center (DC) of the box, speed in the external part (SE) and in the center (SC), and the time spent by mouse in the external part (TE) and the center of the arena (TC). Distance is expressed in arbitrary distance units, speed was expressed as cm/min and time is expressed as a percent of total time (10 min). For open field test, 15 ACN-Ctr and ACN-KO male mice have been used.

#### Hanging wire test

Hanging wire test was performed to assess whole body force and equilibrium (Cordero-Sanchez et al., 2019). The test was conducted in a three-wall box 50 width x 20 depth x 35 cm height, on which, between the tops of side walls, a wire (3 mm diameter) was mounted. Mice were allowed to grasp onto the middle of a wire. Mice attempted to reach one of the wire ends and the number of ‘falls’ or ‘reaches’ was recorded. From an initial score of 10, each fall decreased the score by one and each reach increased the score by 1. The experiment was terminated after 180 s or when the animal reached zero as a score, whichever was earlier. In hanging wire test, 8 ACN-Ctr and 6 ACN-KO male mice were tested.

#### Nesting test

Nesting in small rodents is important for heat conservation, reproduction and shelter, yet it can be a measure of fine coordination and innate memory. For nesting test (Deacon, 2006) mice were housed one per cage. A 5 cm x 5 cm x 1 cm cotton pad was placed in an angle of the cage. After 24 h, the grade of cotton pad shredding was scored from 1 (about 20% of material is shredded) to 5 (all material is shredded and a nest is arranged). In nesting wire test, 11 ACN-Ctr and 11 ACN-KO male mice were tested.

#### Tail suspension test

Tail suspension test, widely used to measure depressive states in small rodents (Can et al., 2012), was conducted in a three wall chamber 55 height x 15 width x 11.5 cm depth. Mouse was suspended to a wire hook by a paper scotch and was recorded for a period of 6 minutes. Escape-like vs immobile behavior time was measured.). 17 ACN-Ctr and 18 ACN-KO male mice were tested.

#### Barnes Maze test

Barnes maze test is used to assess spatial memory and different search strategies in rodents (Gawel et al., 2019; Harrison et al., 2006). The maze was made from a circular, 6-mm thick, white acrylic slab with a diameter of 1 m. Twenty holes with a diameter of 5 cm were made on the perimeter at a distance of 2.5 cm from the edge. This circular platform was then mounted on top of a stool, 60 cm above the ground and balanced. The escape cage was made using a plastic dark box to be easily cleaned. A piece of paper was put in the escape cage, in order to make it more attractive for the mice, and it was changed after every animal test. The maze was placed in the center of a dedicated room and two 120 W lights were placed on the edges of the room facing towards the ceiling about 2 m of the way up from the floor. Four coloured-paper shapes (squares, triangles, circles) were mounted around the room as visual cues, in addition to the asymmetry of the room itself. After testing each mouse, the whole maze was thoroughly cleaned using 70% ethanol and the maze was rotated by 3 holes clockwise to avoid the formation of intra-maze odour cues. All sessions were recorded and videos were analysed with SMART V2.5.21 software. The animals interacted with the Barnes maze in three phases: habituation (Hab), acquisition training (AT), and acquisition probe (AP). After each experimental session the mouse is stored in an holding cage, and after all mice from one home cage completed testing for the day, they were placed back in their home cage together.

On the first day of testing, the habituation phase consisted in placing the mouse in the escape tunnel for 1 min. After that the mouse was placed in the middle of the maze and was allowed to freely explore the maze. The habituation session ended when the mouse entered the escape tunnel or after 5 min elapsed. During the habituation session bight light was turned on but buzzer was turned off. If the mouse did not enter on his own during that time, it was gently nudged to enter and left in the cage for 1 min. The first acquisition training session (AT1) was done 1.5 h after habituation. The mouse was brought in the maze room on an acrylic platform 25×25 cm covered by a cylindrical container. After positioning in the center of the maze and 10 s elapsed the buzzer was switched on, the container was lifted, and the mouse was free to explore the maze. The session ended when the mouse entered the escape tunnel or after 3 min elapsed. If the mouse did not enter the escape box in 3 min, it was gently nudged to enter and left inside for 30 sec. On days 2 (AT2, AT3) and 3 (AT4, AT5) two acquisition trainings per day were performed with 1.5 h of interval between sessions for each animal. On the day 4, three acquisition training sessions were performed (AT6, AT7, AT8) with 15 min of interval between one another, for each animal. On the day 5 acquisition probe (AP) session was performed: the escape cage was removed, the mouse was brought to the center of the maze and, after 10 sec, the buzzer was turned on and the container was removed. Each mouse was given 90 sec to explore the maze, at the end of which, the buzzer was turned off and the mouse was returned to its holding cage.

The search strategies were categorized as follows. The direct (spatial) search was computed when the animal moved directly to the target hole or to an adjacent hole before the target (for the first search) or when mouse approaches target hole from at least 4 holes far away. The random search strategy was computed when at least three visits before the target hole happened in an unsystematic manner, i.e., the animal visited non-adjacent holes and/or crossed the maze. The serial search strategy was computed when there were visits to at least three sequential holes in clockwise or counter-clockwise manner from the target hole. Lastly, return strategy was identified when mouse returned to the target hole from 1-2 adjacent holes. The return strategy can be considered as an exploration of the area around the target hole. In Barnes maze, 10 ACN-Ctr and 10 ACN-KO male mice were tested. During AP session, one ACN-Ctr mouse was excluded because it fell down from the maze and after that showed increased anxiety.

### Electro-encephalographic (EEG) recording

ACN-Ctr and ACN-KO mice were anesthetized with isoflurane (5% (v /v) in 1 L/min O_2_. Four screw electrodes (Bilaney Consultants GMBH, Dusseldorf, Germany) were inserted bilaterally through the skull over the cortex (anteroposterior, +2.0–3.0 mm; left–right 2.0 mm from bregma); a further electrode was placed into the nasal bone as ground. The 5 electrodes were connected to a pedestal (Bilaney, Dusseldorf, Germany) and fixed with acrylic cement (Palavit, New Galetti and Rossi, Milan, Italy). The animals were allowed to recover for a week from surgery before the experiment. EEG activity was recorded in a Faraday chamber using a Power-Lab digital acquisition system (AD Instruments, Bella Vista, Australia; sampling rate 100 Hz, resolution 0.2 Hz) in freely moving awake mice. Basal cerebral activity was recorded continuously for 24 h (from 5 pm to 4 pm). Segments with movement artefacts or electrical noise were excluded from statistical analysis. All EEG traces were also analysed and scored for the presence of spikes, which were discriminated offline with the spike histogram extension of the software (LabChart v8 Pro Windows). Spikes were defined as having a duration < 200 ms with baseline amplitude set to 4.5 times the standard deviation of the EEG signal (determined during inter-spike activity periods, whereas repetitive spiking activity was defined as 3 or more spikes lasting < 5 s)

### Sample size determination

For determination of sample size we avail to the Guidelines for the Care and Use of Mammals in Neuroscience and Behavioral Research (Appendix A, Sample Size Determination) (National Research Council (US) Committee on Guidelines for the Use of Animals in Neuroscience and Behavioral Research, 2003). In particular, for Barnes maze experiment the calculation of sample size for repeated measures was used, while for all other experiments, calculation of sample size for continuous variables was used.

### Statistical analysis

Statistical analysis was performed using GraphPad Prism software v7. For Western blot validation of SG-MS results (Fig. 1C), one sample t-test was used. For comparison of two sample groups, a twotailed unpaired t-test (Fig. 7B; Suppl. Figs. 8B and 9B-D) or unpaired t test with Welch’s correction (Suppl. Fig. 3) were used. For comparison of ACN-KO proteomics with 5xFAD and HE datasets (Fig. 4), hypergeometric test was used. Barnes maze results were analysed as follows: parameters of repeated AT sessions (Fig. 5B) were analysed using two-way ANOVA for repeated measures; single parameters of habituation, AT1 and AP sessions (Figs. 5C,D and 6B) were analysed using a twotailed unpaired t-test; search strategies during AP session (Fig. 5E) were analysed using Chi-square (Fisher’s exact) test; differences in distance and time travelled in external and internal zones of Barnes maze between genotypes (Fig. 6C,D) were analysed by two-way ANOVA Sidak’s multiple comparisons test. Risk to develop seizure between groups was analysed using log-rank (Mantel-Cox) test. qPCR results in Fig. 7C were analysed using one-way ANOVA with Tukey’s posthoc test. The distance, time and speed of mice movements during Open field test (Suppl. Fig. 9A) were analysed using two-way ANOVA Sidak’s multiple comparisons test. In all tests, differences were considered significant at p < 0.05. Detailed report on statistical methods and results is provided in Suppl. Table 9.

## ACKNOWLEDGEMENTS

This work had the following financial supports: grants 2013-0795 to AAG, 2014-1094 to DL from the Fondazione Cariplo; grants FAR-2016 and FAR-2019 to DL from The Università del Piemonte Orientale; L.T. was supported by fellowship from the CRT Foundation (1393-2017); MM received financial support from AGING Project – Department of Excellence – DIMET, Università del Piemonte Orientale.

