## Supplemental Material for "Deletion of calcineurin from astrocytes reproduces proteome signature of Alzheimer’s disease and epilepsy and predisposes to seizures"

[#] These Authors share seniorship.

### Index


| **Content** | **Page** |
|---|---|
| Supplementary Methods. Table 1. **List of oligonucleotide primers used for qPCR** | 2 |
| Supplementary Figure 1. **RNA-seq of ACN-KO vs ACN-Ctr mice** | 4 |
| Supplementary Figure 2. **qPCR validation of RNA-seq and proteomics** | 5 |
| Supplementary Figure 3. **Cell component and Function analysis of DEPs** | 6 |
| Supplementary Figure 4. **OXPHOS, PS, HD and AD-related DEPs** | 7 |
| Supplementary Figure 5. **Putative MAPT-regulated network of HIP DEPs** | 8 |
| Supplementary Figure 6. **Putative PSEN1-regulated network of HIP DEPs** | 9 |
| Supplementary Figure 7. **Putative APP-regulated network of HIP DEPs** | 10 |
| Supplementary Figure 8. **Absence of ultrastructural alterations** | 11 |
| Supplementary Figure 9. **Behavioral analysis of ACN-KO mice** | 12 |
| **Legends for Supplementary Videos 1 and 2** | 13 |

**Supplementary Table 1. List of oligonucleotide primers used in this work.**

| Gene | Accession number | Forward Reverse | Sequence 5' to 3' |
|---|---|---|---|
| S18 | NM_213557 | Forward<br>Reverse | TGCGAGTACTCAACACCAACA<br>CTGCTTTCCTCAACACCACA |
| Calm1 | NM_009790.5 | Forward<br>Reverse | GTCAGAACCCAACAGAAGCC<br>CTCTGGGAAGTCAATGGTGC |
| Col1a1 | NM_007742.4 | Forward<br>Reverse | GTTCAGCTTTGTGGACCTCC<br>TTTCCACGTCTCACCATTGG |
| Cox6a1 | NM_007748.4 | Forward<br>Reverse | CATGCTCAACGTGTTCCTCA<br>ATGAGGGTTGTGGAAGAGGG |
| Glast | NM_148938.3 | Forward<br>Reverse | AATGCCTTCGTTCTGCTCAC<br>ATCCTCATGAGAAGCTCCCC |
| Slc1a2, Glt1 | NM_001077514.3 | Forward<br>Reverse | CTGGTGCAAGCCTGTTTCC<br>TAGTTTCTTCAGGGGCCTCG |
| H1f0 | NM_008197.3 | Forward<br>Reverse | ACCGGTGTTCTCAAGCAAAC<br>CTTGGCTGCCTTCTTTGGAG |
| Atp5j | NM_001302213.1 | Forward<br>Reverse | GGACCTGTTGATATTGGCCC<br>GACTGGGGTTTGTCGATGAC |
| Lmnb1 | NM_010721.2 | Forward<br>Reverse | AGATCGAGCTGGGCAAGTT<br>CTTGATCTGGGCTCCACTGA |
| Slc4a4, Nbce1 | NM_018760.2 | Forward<br>Reverse | GGGGAGGTTGACTTCTTGGA<br>GACTTGGCTTTCCCCTTTGG |
| Pgm1 | NM_025700.3 | Forward<br>Reverse | CCTGACCATCATCCAGACCA<br>CATCGAAGCTGATCACCACG |
| Rps10 | NM_025963.3 | Forward<br>Reverse | GAGACTACCTGCACCTACCC<br>GCCTCCCCTCTTGTGAATCT |
| Serpinab1 | NM_009244.4 | Forward<br>Reverse | CTCAGCCTCGGTCACCAC<br>GAACCATGCAACACAGGCC |
| Thy1 | NM_009382.3 | Forward<br>Reverse | TCGGAACTCTTGGCACCAT<br>CAGGCGAAGGTTTTGGTTCA |
| Tmx2 | NM_025868.4 | Forward<br>Reverse | GCTGTACATGGGTCCTGAGT<br>GGCAAAGAACTCCACAATCCA |
| Mtx2 | NM_016804.4 | Forward<br>Reverse | AGCCGACATGTCTCTGGTG<br>ACAAGGTCGCATTTTCAGGC |
| Rpl6 | NM_011290.5 | Forward<br>Reverse | AGGGGCAAGAGAGTGGTTTT<br>AGAGGTGGCAATGACAAACT |
| Vdac1 | NM_001362693.1 | Forward<br>Reverse | TGGAACACAGACAACACCCT<br>ATGTGCTCCCTCTTGTACCC |
| GFAP | NM_001131020.1 | Forward<br>Reverse | GCTCCAAGATGAAACCAACC<br>GAACTGGATCTCCTCCTCCA |

| | | | |
|---|---|---|---|
| Vapa | NM_001355402.1 | Forward<br>Reverse | TCCTCGACCCTCCTTCAGA<br>CGAGGTGCTGTAGTCTTCAC |
| Rab31 | NM_133685.2 | Forward<br>Reverse | TACGGGAGCTCAAAGTGTGT<br>AAGGCACGGTTTTGGTCATG |
| Iba1 | NM_019467.2 | Forward<br>Reverse | CCGTCCAAACTTGAAGCCTT<br>ACCCCAAGTTTCTCCAGCAT |
| IL-1β | NM_008361.3 | Forward<br>Reverse | AAGTTGACGGACCCCAAAAGA<br>TGTTGATGTGCTGCTGCGA |
| TNFα | NM_013693.2 | Forward<br>Reverse | ACTGAACTTCGGGGTGATCG<br>CTCCTCCACTTGGTGGTTTG |
| Ptgs2,<br>Cox2 | NM_011198.4 | Forward<br>Reverse | GAACAACATCCCCTTCCTGC<br>AGAGGTTGGAGAAGGCTTCC |

A

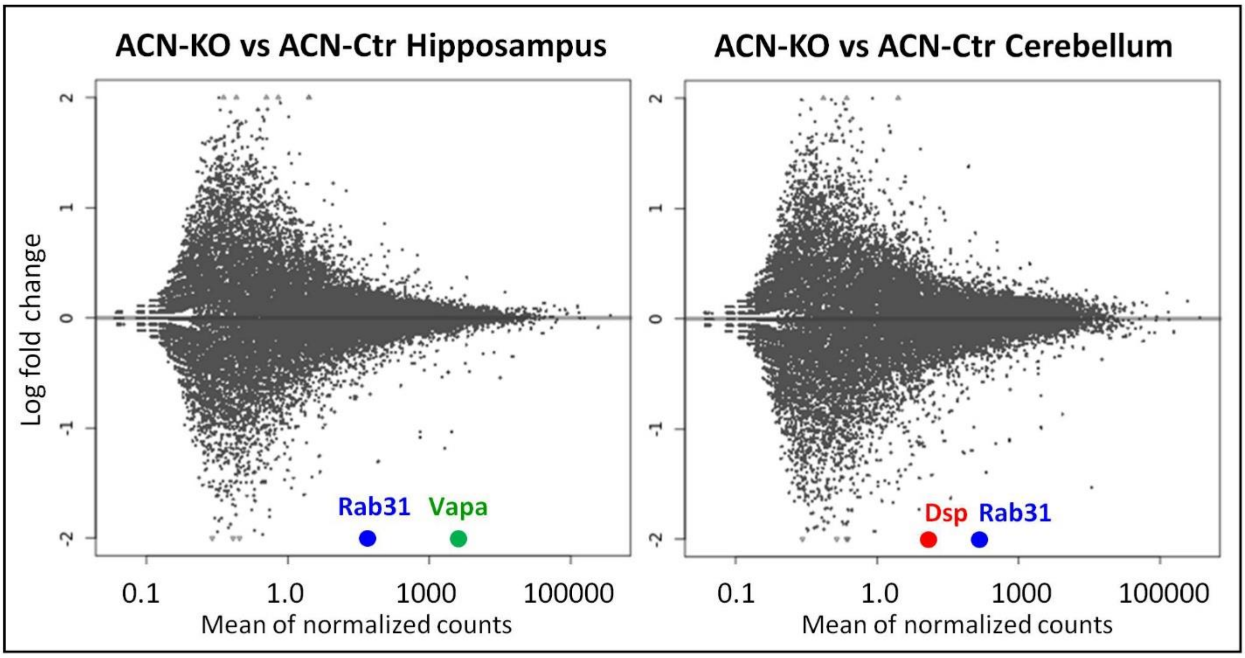


B

| Differentially expressed genes in ACN-KO vs ACN-Ctr mice | | | | |
|---|---|---|---|---|
| **Gene ID** | **Name** | **Expr.lev.** | **FC** | **Adj.p.v.** |
| **Hippocampus** | | | | |
| ENSMUSG00000056515 | **Rab31** | 745.31 | -2.11404 | 8.63E-48 |
| ENSMUSG00000024091 | **Vapa** | 2089.04 | -2.04202 | 4.71E-67 |
| **Cerebellum** | | | | |
| ENSMUSG00000054889 | **Dsp** | 347.93 | -2.62079 | 0.0026 |
| ENSMUSG00000056515 | **Rab31** | 745.31 | -2.14355 | 2.25E-54 |

**Supplementary Figure 1. RNA-seq analysis of hippocampal and cerebellar mRNA of ACN-KO vs ACN-Ctr mice**. RNA-seq analysis was performed using Lexogen (www.Lexogen.com) service. (A) Plot showing distribution of mRNA annotations. Rab31 (blue), Vapa (green) and Dsp (red) were significantly regulated using cut-off of fold change 2 and adjusted p-value < 0.05. (B) There were two differentially expressed genes (DEGs) in both hippocampus (Rab31 and Vapa, both down-regulated respectively, -2.11 and -2.04 fold) and cerebellum (Dsp and Rab31 down-regulated respectively, -2.62, -2.14).

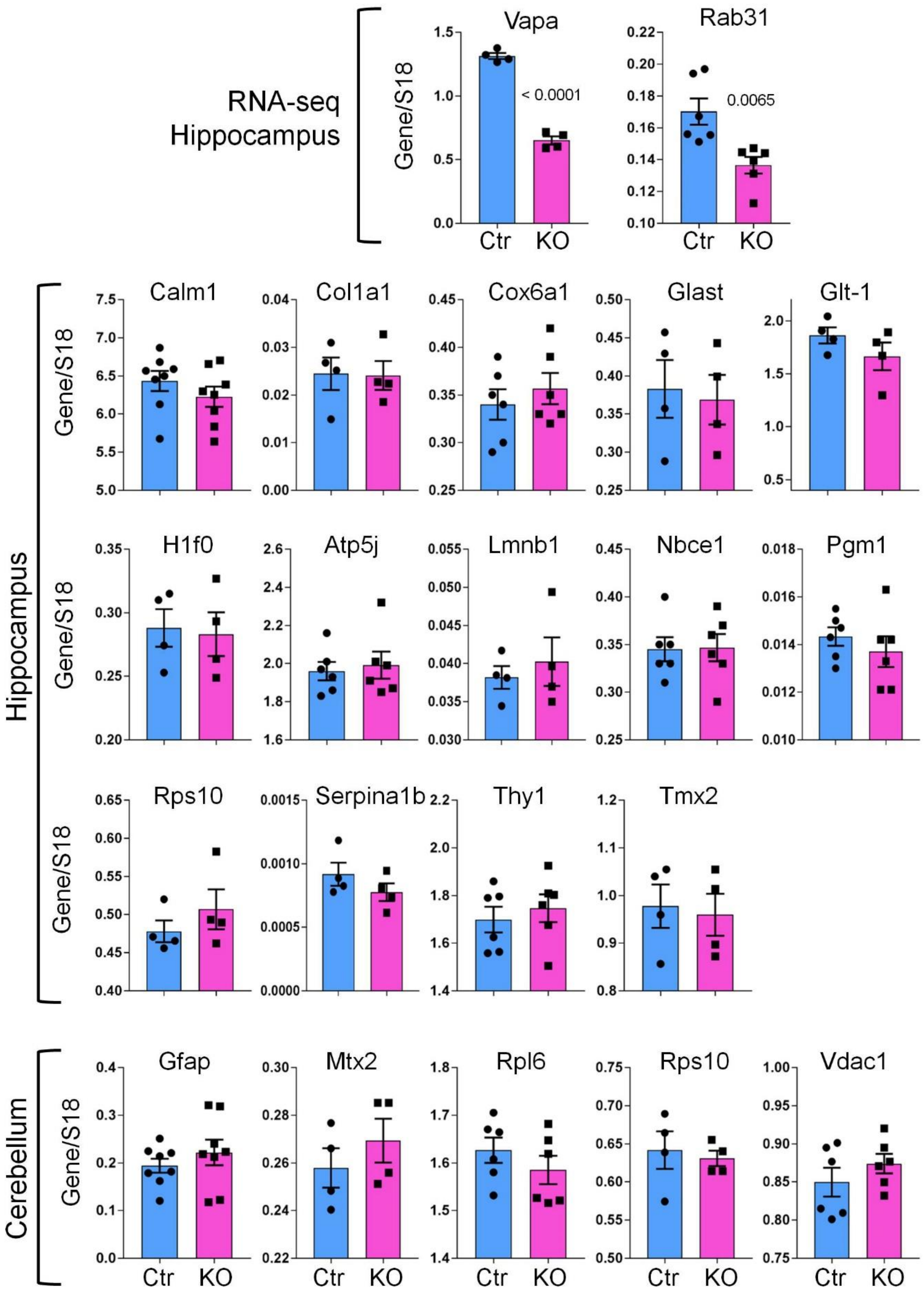


**Supplementary Figure 2. qPCR validation of RNA-seq and shotgun mass spectrometry proteomics**. Note that only Vapa ($p < 0.0001$) and Rab31 ($p = 0.0065$) were significantly down regulated in the hippocampus of ACN-KO vs ACN-Ctr mice.

A. Wht-Hip: Cell component

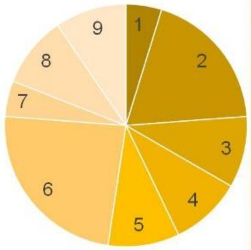


| Series | Cell component | Proteins | % |
|---|---|---|---|
| 1 | Cytoskeleton | 1 | 4.8 |
| 2 | Cytosol | 4 | 19.0 |
| 3 | Endoplasmic reticulum | 2 | 9.5 |
| 4 | Extracellular | 2 | 9.5 |
| 5 | Mitochondria | 2 | 9.5 |
| 6 | Nucleus | 5 | 23.8 |
| 7 | Plasma membrane | 1 | 4.8 |
| 8 | Ribosome | 2 | 9.5 |
| 9 | Vesicles | 2 | 9.5 |

B. Wht-Hip: Function

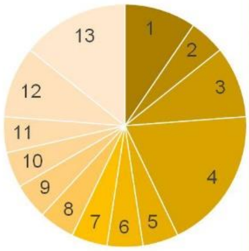


| Series | Function | Proteins | % |
|---|---|---|---|
| 1 | Cell-cell interaction | 2 | 9.5% |
| 2 | Channel | 1 | 4.8% |
| 3 | Chromatin organization | 2 | 9.5% |
| 4 | Metabolism | 4 | 19.0% |
| 5 | Motility | 1 | 4.8% |
| 6 | OxPhos | 1 | 4.8% |
| 7 | RNA processing | 1 | 4.8% |
| 8 | Protein degradation | 1 | 4.8% |
| 9 | Protein processing | 1 | 4.8% |
| 10 | Sctuctural | 1 | 4.8% |
| 11 | Signaling | 1 | 4.8% |
| 12 | Translation | 2 | 9.5% |
| 13 | Vesicle trafficking | 3 | 14.3% |

C. Wht-Cb: Cell component

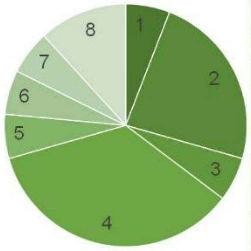


| Series | Cell component | Proteins | % |
|---|---|---|---|
| 1 | Cytoskeleton | 1 | 5.9% |
| 2 | Cytosol | 4 | 23.5% |
| 3 | Endoplasmic reticulum | 1 | 5.9% |
| 4 | Mitochondria | 6 | 35.3% |
| 5 | Nucleus | 1 | 5.9% |
| 6 | Nucleus/cytosol | 1 | 5.9% |
| 7 | Plasma membrane | 1 | 5.9% |
| 8 | Ribosome | 2 | 11.8% |

D. Wht-Cb: Function

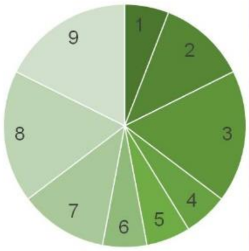


| Series | Function | Proteins | % |
|---|---|---|---|
| 1 | Cell-cell interaction | 1 | 5.9% |
| 2 | Metabolism | 2 | 11.8% |
| 3 | OxPhos | 3 | 17.6% |
| 4 | Protein processing | 1 | 5.9% |
| 5 | RNA processing | 1 | 5.9% |
| 6 | Scaffold | 1 | 5.9% |
| 7 | Signaling | 2 | 11.8% |
| 8 | Translation | 3 | 17.6% |
| 9 | Transport | 3 | 17.6% |

E. Syn-Hip: Cell component

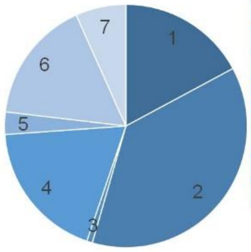


| Series | Cell component | Proteins | % |
|---|---|---|---|
| 1 | Cytoskeleton | 23 | 17.2% |
| 2 | Cytosol | 50 | 37.3% |
| 3 | Golgi apparatus | 1 | 0.7% |
| 4 | Mitochondria | 25 | 18.7% |
| 5 | Nucleus | 4 | 3.0% |
| 6 | Plasma membrane | 22 | 16.4% |
| 7 | Vesicle | 9 | 6.7% |

F. Syn-Hip: Function

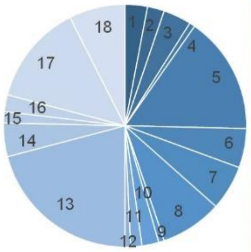


| Series | Function | Proteins | % |
|---|---|---|---|
| 1 | Cell-cell interaction | 4 | 3.0% |
| 2 | Chaperone | 3 | 2.2% |
| 3 | Exocytosis | 5 | 3.7% |
| 4 | Homeostasis | 1 | 0.7% |
| 5 | Metabolism | 21 | 15.7% |
| 6 | Microtubules | 7 | 5.2% |
| 7 | Motility | 8 | 6.0% |
| 8 | OxPhos | 11 | 8.2% |
| 9 | Pore complex | 1 | 0.7% |
| 10 | Protein degradation | 3 | 2.2% |
| 11 | Protein processing | 2 | 1.5% |
| 12 | Scaffold | 1 | 0.7% |
| 13 | Signaling | 28 | 20.9% |
| 14 | Structural | 6 | 4.5% |
| 15 | Transcription | 2 | 1.5% |
| 16 | Translation | 3 | 2.2% |
| 17 | Transport across membrane | 18 | 13.4% |
| 18 | Vesicle trafficking | 10 | 7.5% |

G. Syn-Cb: Cell component

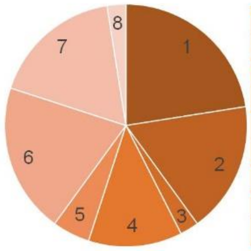


| Series | Cell component | Proteins | % |
|---|---|---|---|
| 1 | Cytoskeleton | 9 | 22.5% |
| 2 | Cytosol | 7 | 17.5% |
| 3 | Golgi apparatus | 1 | 2.5% |
| 4 | Mitochondria | 5 | 12.5% |
| 5 | Nucleus | 2 | 5.0% |
| 6 | Plasma membrane | 8 | 20.0% |
| 7 | Ribosome | 7 | 17.5% |
| 8 | Vesicles | 1 | 2.5% |

H. Syn-Cb: Function

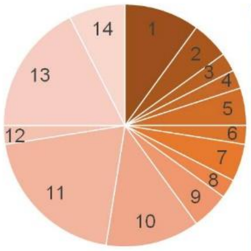


| Series | Cell component | Proteins | % |
|---|---|---|---|
| 1 | Cell-cell interaction | 4 | 10.0% |
| 2 | Channel | 2 | 5.0% |
| 3 | Chaperone | 1 | 2.5% |
| 4 | Immune response | 1 | 2.5% |
| 5 | Metabolism | 2 | 5.0% |
| 6 | Motility | 1 | 2.5% |
| 7 | OxPhos | 2 | 5.0% |
| 8 | Protein processing | 1 | 2.5% |
| 9 | Protein trafficking | 2 | 5.0% |
| 10 | Signaling | 5 | 12.5% |
| 11 | Structural | 8 | 20.0% |
| 12 | Transcription | 1 | 2.5% |
| 13 | Translation | 7 | 17.5% |
| 14 | Transport across membrane | 3 | 7.5% |

**Supplementary Figure 3. Cell component and Function analysis.** Cell component and Function analysis of differentially regulated proteins. in whole-tissue hippocampal (Wht-Hip) (A, B) and cerebellar (Wht-Cb) (C, D) tissues, and in hippocampal (Syn-Hip) (E, F) and cerebellar (Syn-Cb) (G, H) synaptosomal fractions.

Common proteins in OxPhos, PD, HD, and AD pathways

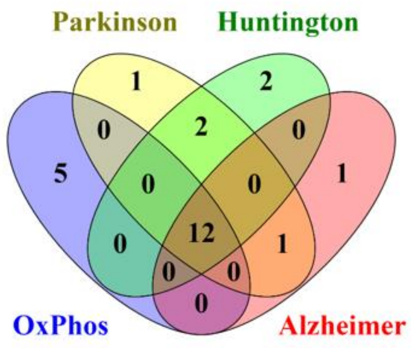


| No | Uniprot ID | Protein | p.v. | FC | Mito com |
|---|---|---|---|---|---|
| 1 | NDUBA_MOUSE | NADH dehydrogenase [ubiquinone] 1 beta subcomplex subunit 10 | 0.0298 | 9.80 | Com I |
| 2 | CX6A1_MOUSE | Cytochrome c oxidase subunit 6A1 | 0.0173 | 4.17 | Com IV |
| 3 | ATPB_MOUSE | ATP synthase subunit beta | 0.0004 | 1.84 | Com V |
| 4 | ATPA_MOUSE | ATP synthase subunit alpha | 0.0147 | -1.28 | Com V |
| 5 | UCRI_MOUSE | Cytochrome b-c1 complex subunit Rieske | 0.0229 | -1.86 | Com III |
| 6 | CX7A2_MOUSE | Cytochrome c oxidase subunit 7A2 | 0.0275 | -2.85 | Com IV |
| 7 | ATP8_MOUSE | ATP synthase protein 8 | 0.0372 | -3.15 | Com V |
| 8 | ATPD_MOUSE | ATP synthase subunit delta | 0.0053 | -3.54 | Com V |
| 9 | ATPO_MOUSE | ATP synthase subunit O | 0.00004 | -3.81 | Com V |
| 10 | NDUA4_MOUSE | Cytochrome c oxidase subunit NDUFA4 | 0.0061 | -3.88 | Com I |
| 11 | AT5G1_MOUSE | ATP synthase F(0) complex subunit C1 | 0.0003 | -9.63 | Com V |
| 12 | ATP5J_MOUSE | ATP synthase-coupling factor 6 | 0.0080 | -11.46 | Com V |

**Supplementary Figure 4. Intersection of overrepresented KEGG pathways revealed presence of 12 common proteins, components of mitochondrial OXPHOS pathway.** Hippocampal DEPs were subjected to gene ontology analysis using DAVID tool. Intersection of KEGG pathways Parkinson's disease, Huntington's disease, Alzheimer's disease and OXPHOS revealed presence of 12 common proteins, listed in the table below.

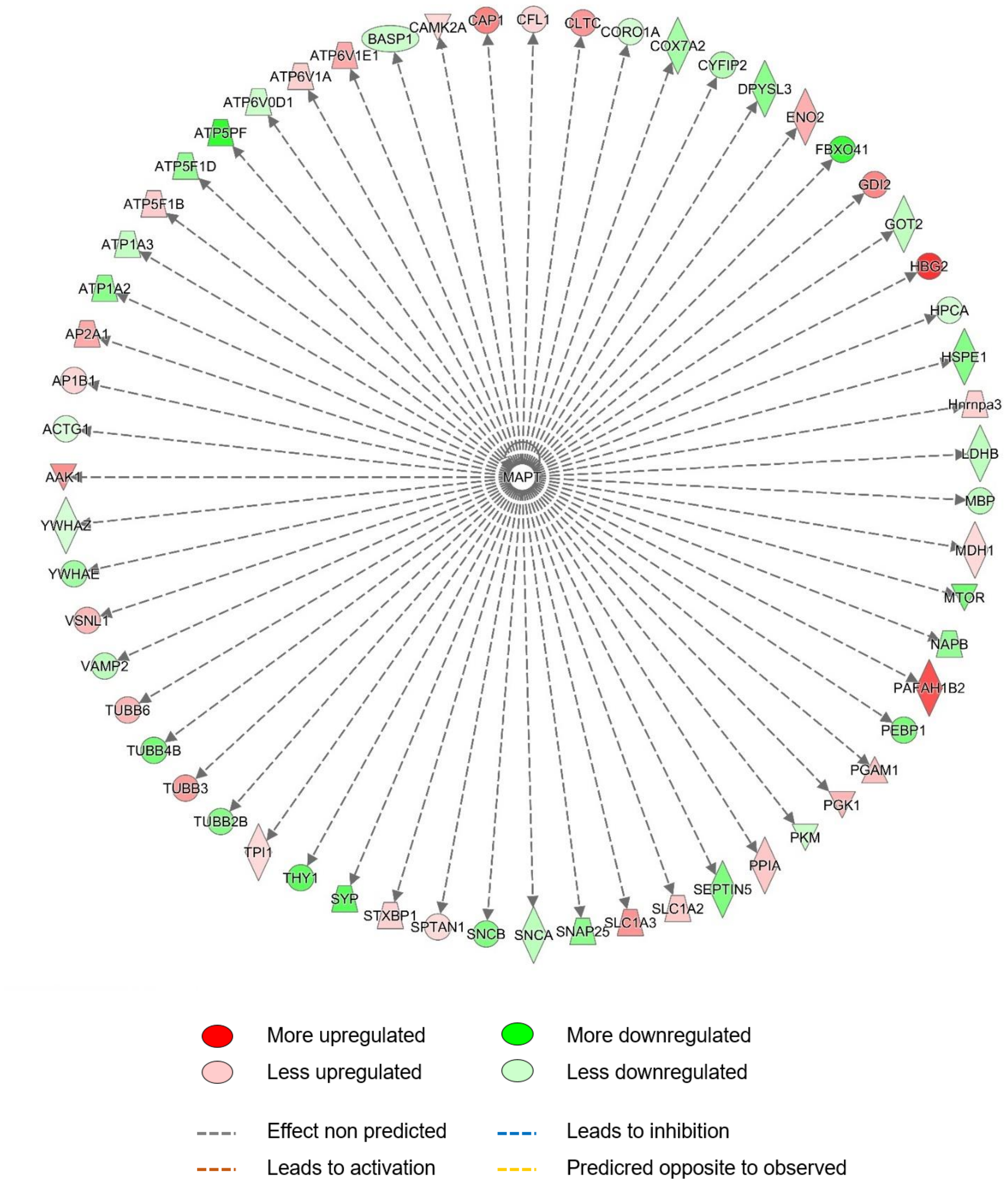


**Supplementary Figure 5.** MAPT-regulated network identified by IPA Upstream Regulator analysis.

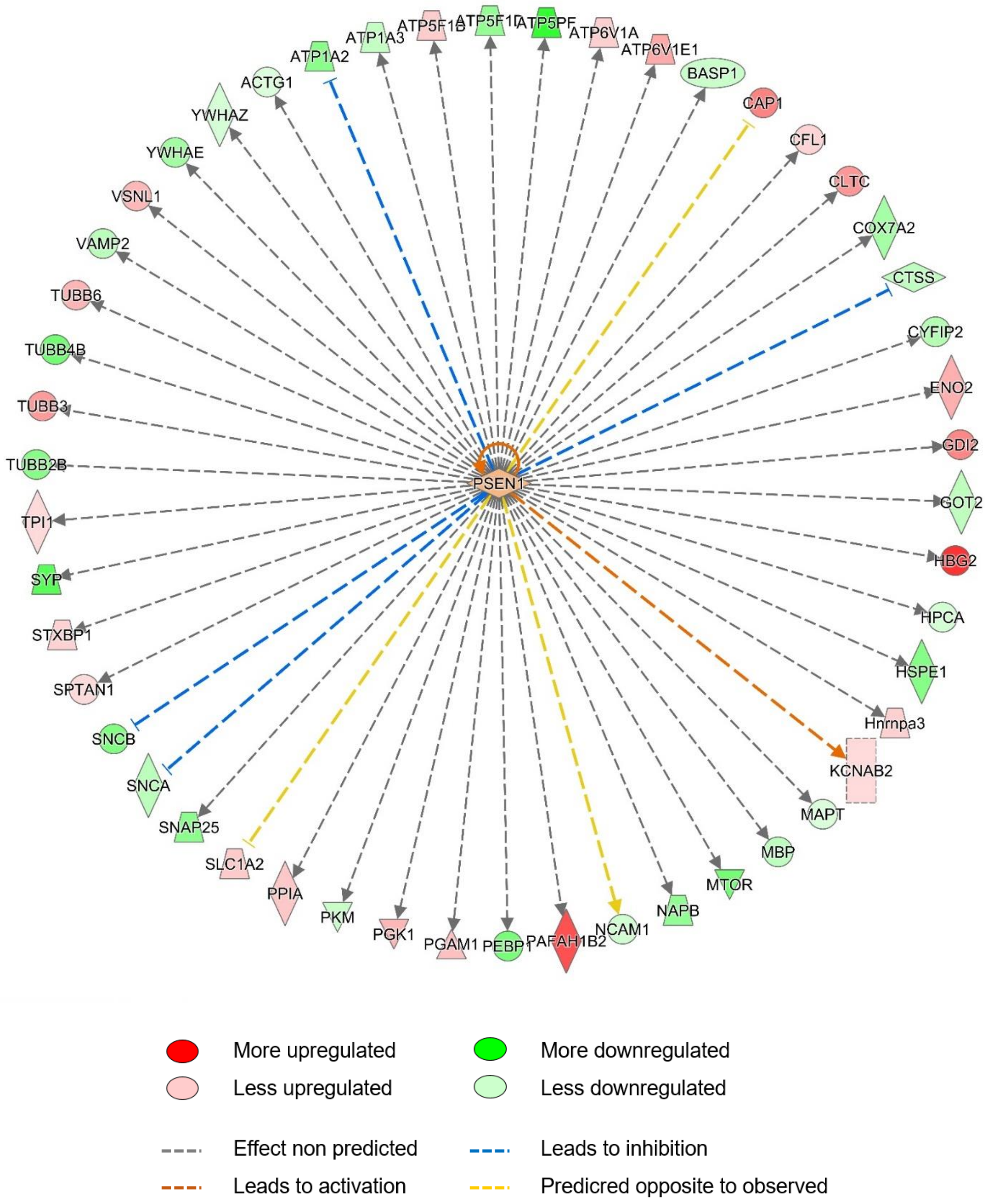


**Supplementary Figure 6.** PSEN1-regulated network identified by IPA Upstream Regulator analysis.

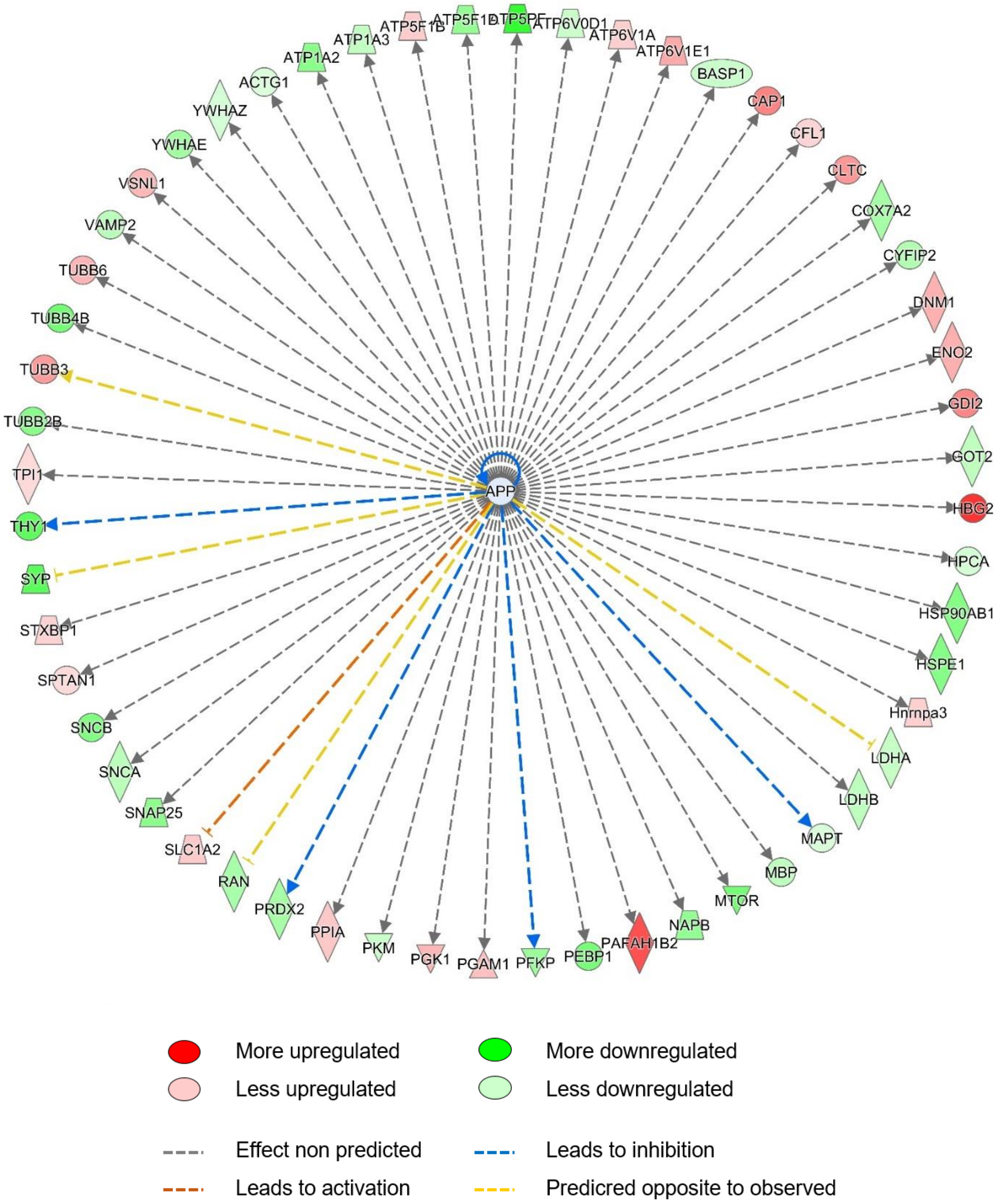


**Supplementary Figure 7.** APP-regulated network identified by IPA Upstream Regulator analysis.

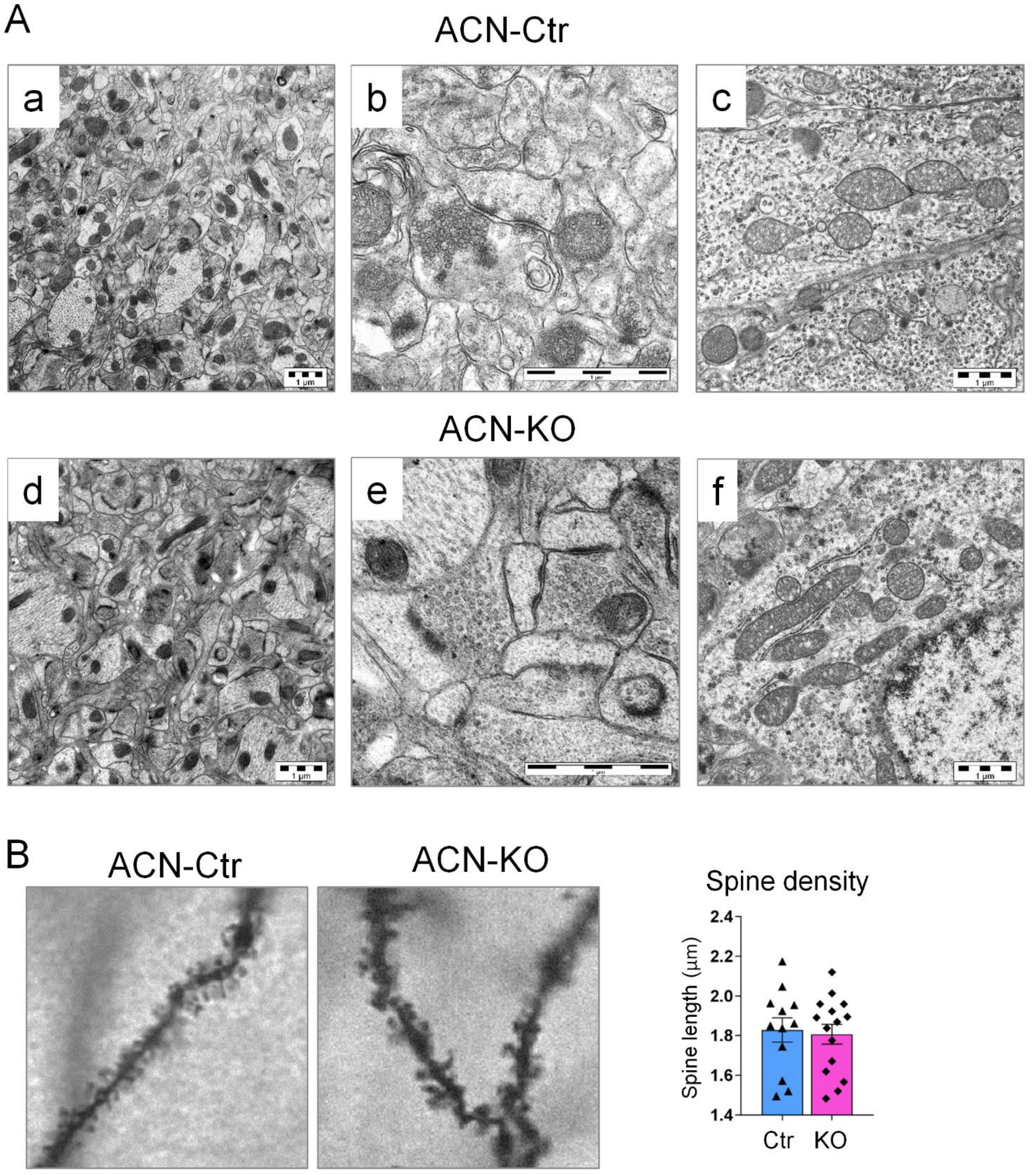


**Supplementary Figure 8. Absence of ultrastructural alterations.** (A) Transmission electron microscopy images of CA1 hippocampal region of ACN-Ctr and ACN-KO mice. Note absence of abnormalities in general neuropil organization (a and d), in synaptic structure (b and e) and in mitochondria (c and f). Bar, 1 µm. (B) Golgi stain images and dendritic spine quantification shows no differences in density of dendritic spines between ACN-Ctr and ACN-KO mice (n = 12 neurons from 4 mice for ACN-Ctr, n = 15 from 5 mice for ACN-KO, p = 0.78).

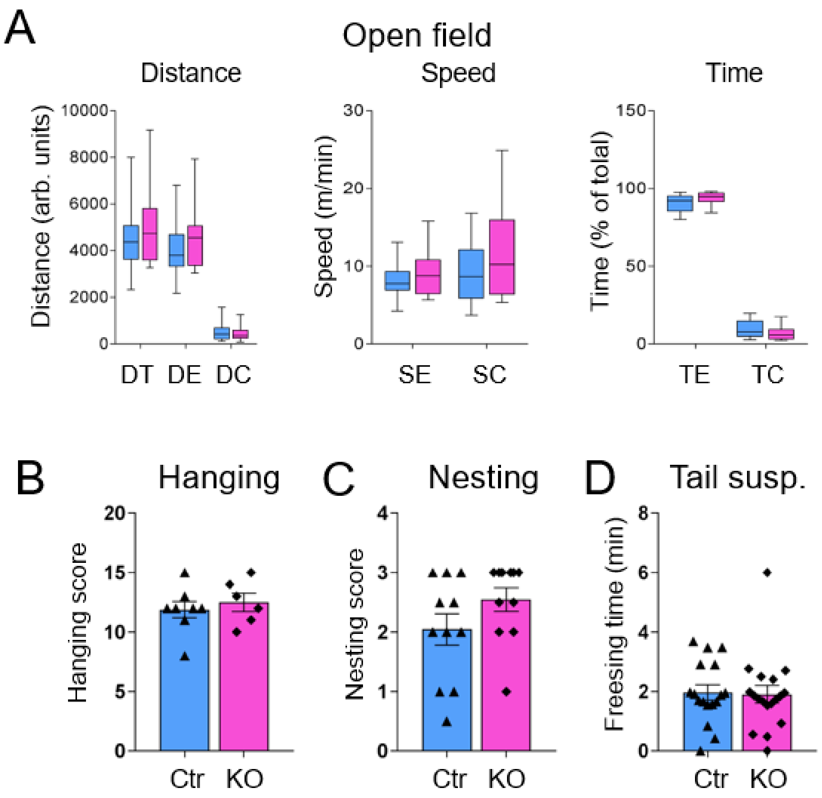


**Supplementary Figure 9. Normal motor behaviour, absence of anxiety and depression in ACN-KO mice.** (A) Open field test shows no alterations in general motor and explorative activity and anxiety in ACN-KO (n = 15) compare to ACN-Ctr mice (n = 15) (DT, total distance, p = 0.73; DE, distance travelled in external part of arena, p = 0.52; DC, distance travelled in central part, p = 0.98; SE, speed in external part, p = 0.72; SC, speed in central part, p = 0.25; TE, time spent in external part, p = 0.15; TC, time spent in central part, p = 0.35). (B) Hanging test (p = 0.56) shows no alterations in body force and equilibrium in ACN-KO (n = 6) vs ACN-Ctr (n = 8). (C) No differences in nesting behaviour (n = 11, p = 0.145). (D) Normal depressive state (p = 0.86) in ACN-KO (n = 18) compare with ACN-Ctr (n = 17). (E) Novel object recognition test (NORT) revealed a trend to a lower discrimination memory index in ACN-KO (n = 21) compare with ACN-Ctr mice (n = 16, p = 0.055).

**Legends to Supplementary Videos:**

**Supplementary Video 1. Seizure in 7.8 month-old ACN-KO female**. Note that seizure starts upon bothering of the mouse by slightly pulling the tail. The intensity of seizures corresponds to stage 5 (rearing and falling with forelimb clonus) according to Racine scale.

**Supplementary Video 2. Seizure in 6.7 month-old ACN-KO male**. Note that seizure starts with stage 4 according to Racine scale (forelimb clonus) and develops to stage 7 according Pinel and Rovner scale (jumping).
